# Overlapping MHC class I/II Epitopes Program cDC1-like Differentiation of Monocyte-Derived Dendritic Cells via mTORC1 Signaling Inhibition

**DOI:** 10.64898/2026.04.06.716309

**Authors:** SB Amanya, A Murthy, N Bisht, ZN Bullock, KJ Ernste, J Vasquez-Perez, W Liu, A Trivedi, D Oyewole-Said, A Paul, L Umokoro, V Akhanov, M Samuel, Z Shi, MH Nguyen, S Huang, M Jeong, P Iakova, KA Jain, K Pham, DC Kraushaar, V Konduri, WK Decker

## Abstract

Viral infection polarizes monocyte-derived dendritic cells (moDC) to initiate type 1 immunity. The availability of overlapping (homologous) MHC class I and II epitopes, an occurrence frequently and primarily associated with intracellular infection, significantly enhances this process; however, the underlying mechanism(s) are unclear. We demonstrate that moDC loaded with homologous MHC epitopes acquire a cDC1-like phenotype in a process governed by mTORC1. mTORC1 pathway inhibition leads to NF-κB-mediated expression of IL-12 and other type I immune polarizing genes. The observed cDC1-like gene signature was also significantly enhanced in clinical moDC vaccine products made through methodologies that enforced class I and II antigenic homology. Collectively, these findings reveal a novel and previously unrecognized mechanism of immune governance that might also be exploited in cancer immunotherapy.

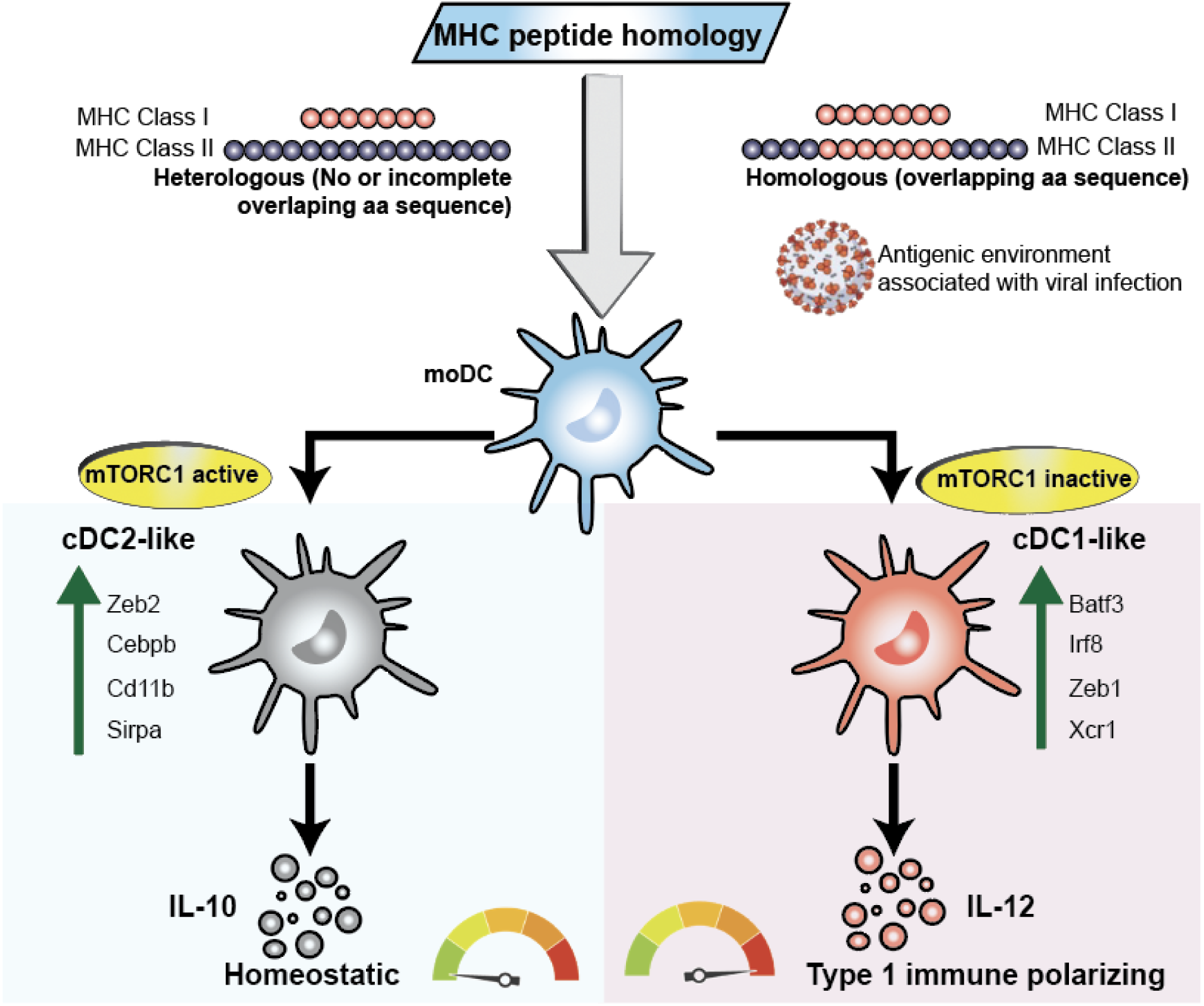

## Background

Monocytes are multipotent cells of the peripheral immune system that play a critical role in immune homeostasis, acting as early responders to acute inflammation(*1, 2*). Inflammatory cues rapidly recruit monocytes to inflamed tissues where they differentiate into specialized antigen presenting cell (APC) populations, depending on prevailing environmental cues. Local GM-CSF and IL-4 are such cues that induce the differentiation of monocyte-derived dendritic cells (moDC), which support tissue resident conventional dendritic cells (cDC) in mounting robust adaptive immune responses as appropriate(*2, 3*).

Viral infection is an immune perturbation which drives moDCs to generate type 1 immune polarization(*4–6*), providing a blueprint for differentiating moDC into inducers of type 1 immune responses that may be effective against tumors. During infection, viruses infect both moDCs and the surrounding tissue, an environment that generates a rich pool of intracellular and extracellular MHC class I and II epitopes, the amino acid sequences of which share substantial sequence overlap(*7–9*). Indeed, immunodominant MHC peptide epitopes with complete MHC class I and II sequence overlap have been reported in various viral infections including lymphocytic choriomeningitis virus (LCMV) infection(*8*), hepatitis C virus infection in which overlapping epitopes exhibited the highest binding affinities(*10*), and influenza virus infection in which viral proteins that generate overlapping MHC epitopes were the most efficiently presented and thus led to the highest levels of CD4 and CD8 T cell activation (*7*). The Yellow fever (YF-17DD) vaccine, one of the safest and most effective of the live attenuated vaccines, also generates overlapping class I and class II epitopes which are highly promiscuous and have been shown to specifically promote the development of circulating memory T cells(*11*). Additionally, overlapping MHC peptides derived from the human glutamic decarboxylase (GAD) 65 protein have been reported to directly drive cytotoxic T-cell responses associated with insulin-dependent diabetes mellitus (IDDM)(*12*). Collectively, these reports position overlapping MHC class I and II epitopes as a powerful antigenic pattern associated with the elicitation of type 1 immune responses, which need not necessarily always be viral in origin.

Leveraging these insights, our group has conscripted this mechanism, a strategy known as homologous antigenic loading, to induce moDC-mediated type 1 immune responses against nonviral tumor targets. In this strategy, moDCs are loaded with antigens that enforce the generation of homologous MHC epitopes using two complementary approaches: loading autologous tumor-derived mRNA and cell lysates to promote MHC class I/II epitope homology, or loading of MHC binding peptides that share identical stretches of amino acid sequencing (i.e. overlapping) between MHC class I and MHC class II(*13, 14*). Either approach generates DC of an identical phenotype, characterized by increased IL-12 production, highly upregulated IFN-_γ_ production from reactive T-cells, and generation of antigen-specific CD8^+^ T-cells with enhanced cytotoxic capabilities(*13–17*). Yet, despite documentation of substantially enhanced type I immune outputs, the mechanism(s) through which homologous MHC epitopes reprogram moDCs into a more potent, type 1 polarizing state remain poorly defined.

Utilizing single cell RNA sequencing (scRNA-seq), we found that mouse BMDCs loaded with homologous MHC epitopes upregulated a cDC1-like cluster characterized by high expression of *Batf3, Irf8, Zeb1, Zfp366* and *Ly75,* and high enrichment in antiviral response pathways. Through transcriptomic and proteomic profiling of both mouse BMDC and human moDC, we linked the induction of the cDC1-like phenotype to mTORC1 inhibition, leading to NF-κB-mediated induction of IL-12 gene expression. Inhibition of mTOR with rapamycin enhanced the cDC1-like phenotype as well as DC-mediated differentiation of cytotoxic T-cell responses. Importantly, the cDC1-like gene signature characterized in the mouse BMDC, was also upregulated in human clinical moDC vaccine products generated through a methodology that enforces homologous class I and II antigenic loading, demonstrating a phenotype that’s highly conserved throughout evolution and with clinical translational potential. This work suggests that homologous MHC epitopes serve as a recognizable pattern, the presence of which generates a cDC1-like phenotype through inhibition of mTORC1 pathway signaling.

## Results

### Loading of BMDCs with homologous MHC epitopes skews differentiation towards a cDC1-like phenotype

In previous work, we have shown that moDCs loaded with homologous antigens heavily skew downstream T-cell responses toward type 1 immunity(*13–17*). To better characterize this homologous MHC epitope-induced phenotype, scRNA-seq was performed on bone marrow-derived DCs (BMDCs) loaded with the perfectly homologous LCMV glycoprotein MHC binding pair KAVYNFATC and GI**<u>KAVYNFATC</u>**GIFA or the heterologous LCMV glycoprotein binding pair KAVYNFATC and DIY**<u>K</u>**G**<u>VY</u>**Q**<u>F</u>**KSVEFD and matured for 24 hours (**Figure 1A**). At this post-maturation timepoint, unsupervised seurat(*18*) clustering of scRNA-seq data revealed 10 transcriptionally distinct Uniform Manifold Approximation and Projection (UMAP) clusters (**Figure 1B**). Of these, clusters 0, 1, and 2 exhibited substantial statistical differences between the peptide sequence overlapping(homologous) and peptide sequence non-overlapping (heterologous control) groups (**Figure 1C** and **Table S1A**). Cluster 0 was enriched for cDC2-associated markers including *Cebpb, Itgam, Cd302, Clec4d, Clec4a2,* and *Cd300c2,* while cluster 1 exhibited strong expression of cDC1-associated markers including *Batf3, Irf8, Irf5, Apol7c, Serpin6b, and Ly75* (**Figure 1D, S1B**). Other minor clusters shared characteristics with either cluster 1 (3 and 9), or cluster 0 (2, 5, and 8), or both 0 and 1 (4, 6, and 7), potentially representing transitional states to the 0 and 1 terminally differentiated clusters (**Figure 1D, Figure S1B**). To discern differentiation state, we performed differentiation trajectory inference using Monocle(*19*). In this analysis, we used cluster 6 as the root based on high expression of *Mafb* (**Figure S1C**), an early transcription factor in myeloid progenitors whose inhibition by PU.1 is necessary to initiate dendritic cell fate commitment(*20*). Additionally, the expression patterns of *Id2* and *Zeb2* pointed to cluster 6 as the root, with *Id2* expression arising from cluster 6 and trending towards cluster 1 while *Zeb2* trended towards cluster 0 (**Figure S1C**). This analysis revealed a branching trajectory with two terminal differentiation states at Cluster 1 and cluster 0 (**Figure 1E**). Pseudotime progression along cluster 0 differentiation branch was associated with high expression of *Fth1, Arg1, Chil3, Ccl9,* and *Clec4d* genes, while positioning along cluster 1 pseudotime branching trajectory was associated with *Cst3, Ccr7, Cd86, Serpinb6b,* and *H2-Aa* expression (**Figure 1F, S1D, S1F**). Additionally, cDC1 canonical transcription factors *Irf8, Batf3,* and *Zfp366* were associated with pseudotime along the cDC1-like cluster differentiation trajectory while *Zeb2, Cd14,* and *Itgam* were associated with the cDC2-like cluster 0 differentiation trajectory (**Figure S1E**). These findings point to cluster 0 and cluster 1 as terminal states with other clusters as transitory states. Because cluster 0 and 1 are terminal states with cell population size that’s significantly different between the MHC binding peptide groups, we focused on these for further analysis. In-depth characterization showed that cluster 1 expressed high levels of cDC1 defining transcription factors and markers including *Irf8, Batf3, Zfp366, Nfil3, Zeb1,* and *Ly75* while cluster 0 was enriched for cDC2 transcription factors and markers that included *Cebpb, Zeb2, Itgam, Sirpa, Cd24a, and Clec4d* (**Figure 1G**). In addition to its cDC1 defining features, cluster 1 exhibited high expression of costimulatory molecule genes including *Cd80, Cd86, Cd83, Cd40, Tnfsf4* (OX40L), and *Tnfsf11*(RANKL); antigen processing and presentation genes: *Tap1, Tap2, B2m, Apol7c, Psmb8, Psmb9*; granzyme inhibitors: *Serpinb6b, Serpinb9, Serpinb6a, and Serpinb9b*, and cell adhesion and migratory genes: *Ccr7, Cd48, Icam1,* and *Ccl17* (**Figure 1H**). These findings suggest that the loading of BMDCs with homologous MHC epitopes induces a transcriptional state closely analogous to that of cDC1, the DC subset most highly specialized in the generation of CD8^+^ T-cell immunity. To validate these findings, we used flow cytometry to characterize the expression of lineage defining markers for cDC1 (Xcr1) and cDC2 (CD11b). As expected, we found high expression of Xcr1 (**Figure 1I**) among DCs loaded with homologous peptides in comparison to controls loaded with heterologous peptides. In contrast, CD11b expression was reduced among BMDCs loaded with homologous epitopes in comparison to heterologous control (**Figure 1J**). Collectively, these findings support a model by which loading of BMDCs with homologous MHC epitopes induces a cDC1-like cell fate.

**Figure 1.**
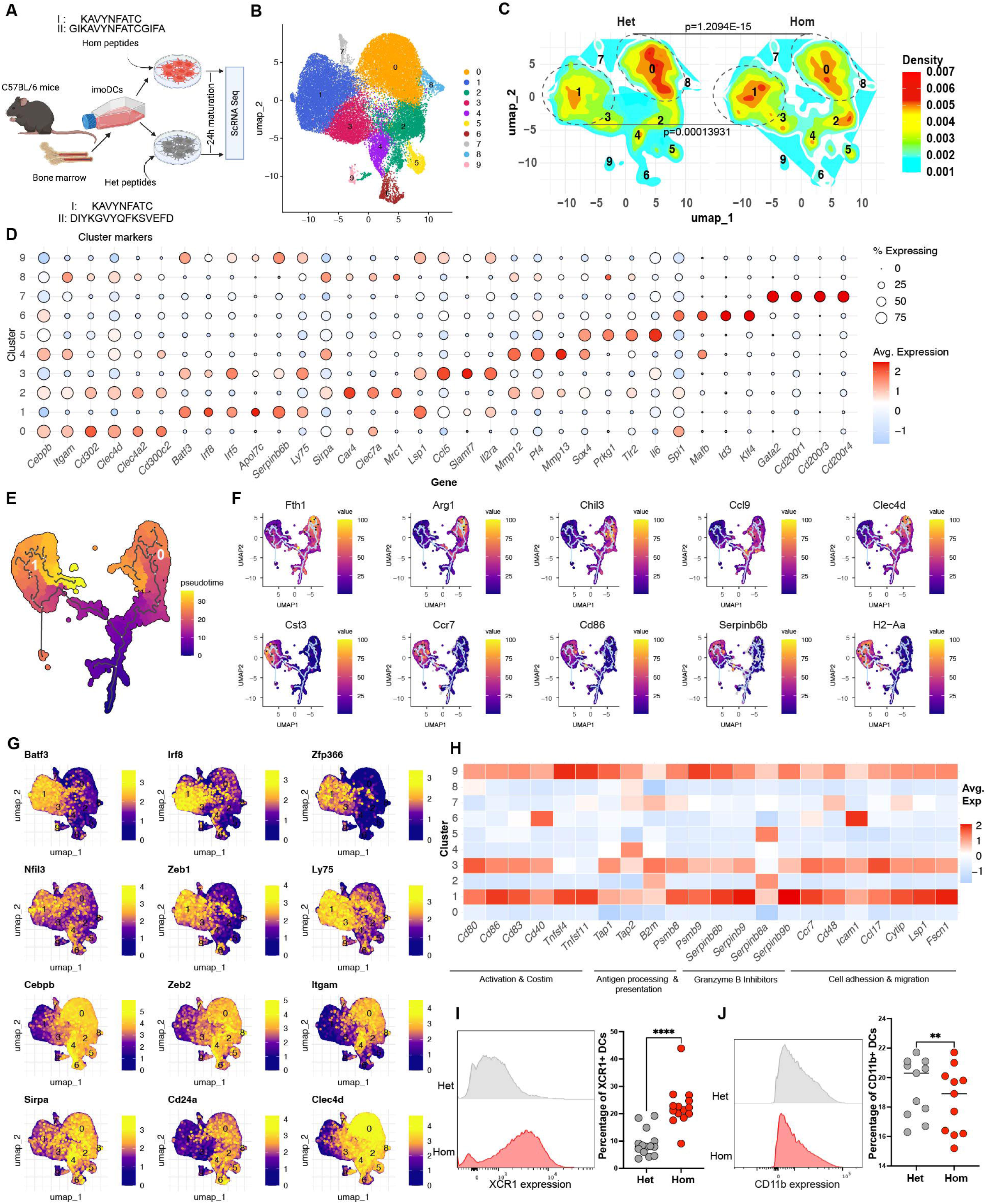
Loading of BMDCs with homologous MHC epitopes skews differentiation towards a cDC1-like phenotype. **A.** Experimental strategy for scRNA-seq analysis of mouse BMDC. Immature BMDC were pulsed with homologous (overlapping) or heterologous (non-overlapping) MHC epitopes and matured for 24 hours prior to harvesting for analysis by scRNA-seq. Three biological replicates were included per peptide loading condition. **B.** UMAP of pooled scRNA-seq data showing 10 transcriptionally distinct clusters of BMDC identified by unsupervised clustering. **C.** UMAP density heatmap demonstrating differences in cluster enrichment between peptide loading groups. Statistical significance of cluster distribution between groups was determined by chi-squared analysis. **D.** Dot plot of cluster-defining marker genes across identified BMDC clusters. Dot size indicates percentage of cells expressing the gene, color intensity represents scaled gene expression levels (red, highest gene expression and blue, lowest gene expression). **E.** Single cell trajectory inference performed by Monocle, projected onto UMAP space to visualize differentiation relationships between clusters. Cluster 6 was designated as the root. **F.** UMAP feature plots showing genes associated with pseudotime progression toward cluster 0 or cluster 1. **G.** Feature plot showing expression of lineage-defining transcription factors and marker genes. Color indicates gene expression level (yellow, highest expression; purple, lowest expression). **H.** Heatmap showing differential expression of functional gene modules associated with DC activation and costimulation, migration, antigen presentation, and granzyme inhibitors across clusters. Color indicates gene expression levels (red, highest expression; blue lowest expression). **I.** Representative histogram overlay and quantification of flow cytometry data showing Xcr1 expression in BMDC under indicated conditions. N=14 biological replicates, statistical analysis performed by paired two-tailed t-test. ****P<0.0001, **P<0.01. **J.** Representative histogram overlay and quantification of flow cytometry data showing CD11b expression in BMDC under the indicated conditions. N=11 biological replicates, statistical analysis performed by paired two-tailed t-test. ****P<0.0001, **P<0.01.

### A type 1 immune polarizing gene program is enriched within cDC1-like cluster 1

The hallmark of cDC1 is an ability to activate cytotoxic CD8^+^ T-cell responses through a coordinated network of type 1 immune polarizing programs(*21, 22*). Compared to the cDC2-like cluster 0, the cDC1-like cluster 1 exhibited high expression of genes related to type 1 immune activation(*21, 23*) including *Ccl22, Fscn1, Ccl17, Socs2, Zeb1, Ktn1, Irf5, Stat2, Stat4, Traf1, Relb, Camk4,* and *Apobec3* (**Figure 2A**), supporting the acquisition of a type 1 immune polarizing phenotype. Consistent with this observation, KEGG analysis revealed enrichment of cluster 1 with transcriptional pathways related to viral infection (**Figure 2B**). To examine signaling pathways underlying these anti-viral response programs, we used MsigDB-curated gene sets(*24*) to characterize specific signaling pathways driving anti-viral response transcriptional programs. Gene Set Enrichment Analysis (GSEA) revealed a positive enrichment of antigen processing and presentation pathway related genes in cluster 1 (NES=1.2, **Figure 2C**), with strong expression of *H2-Eb1, H2-Aa, Cd74, H2-Ob, Hfe, Relb, Psmb9, Psmb8, Psme1, & Psme2* genes driving this positive enrichment (**Figure 2D**). Similarly, the IL-12 signaling pathway, known to direct cDC1-mediated CD8^+^ T-cell activation(*25, 26*), was significantly enriched in cluster 1 (NES=1.52, **Figure 2E**). This positive enrichment was primarily driven by strong expression of *Cst3*, *Cd40, Gnb4, Pkib, Irf8, Dusp5, Slc33a1, Ptpn1, Samhd1, Basp1, Ifi35, Stat3, Stat2, Tspo, Zyx,* and *Stat1* (**Figure 2F**). The IL-2 signaling pathway, known to increase DC expansion and activation(*27*), was also significantly enriched in the cDC1-like cluster 1 with NES=1.48 (**Figure 2G**). The genes driving this positive enrichment included *Cst3, Serpinb9, Cd40, Gnb4, Apol7c, Pkib, Ccnd2, Syngr2, Irf5, Psmb9, Psmb10, Fnbp11, Marcks,* and *Irf8* (**Figure 2H**). Other upregulated pathways included antigen processing and proteosomal degradation pathways (**Figure S2A/B**), antigen processing and presentation on MHC II (**Figure S2C/D**), and IL-2/Stat5 signaling (**Figure S2D/E**). In contrast to the pathways listed above, the anti-inflammatory IL-10 pathway was significantly downregulated in cluster 1 (NES= ^-^1.91, **Figure 2I**) with antiinflammatory genes including *Socs3, Lilrb4b, Srgn, Tpd52, Cstd, Ctsl, Ccr1, Saa3, Wfdc17, Vsir, F10, Tgfb1*, and *Hp* driving the negative enrichment of the IL-10 pathway (**Figure 2J**). Additionally, the anti-inflammatory IL-6-Jak-Stat3 signaling pathway was also downregulated in cluster 1 compared to cluster 0 (**Figure S2F/G**). These findings reveal an extensive transcriptional rewiring program through which a positive enrichment of type 1 immune polarizing pathways and negative enrichment of anti-inflammatory pathways establish the anti-viral transcriptional program seen in the cDC1-like cluster 1.

**Figure 2.**
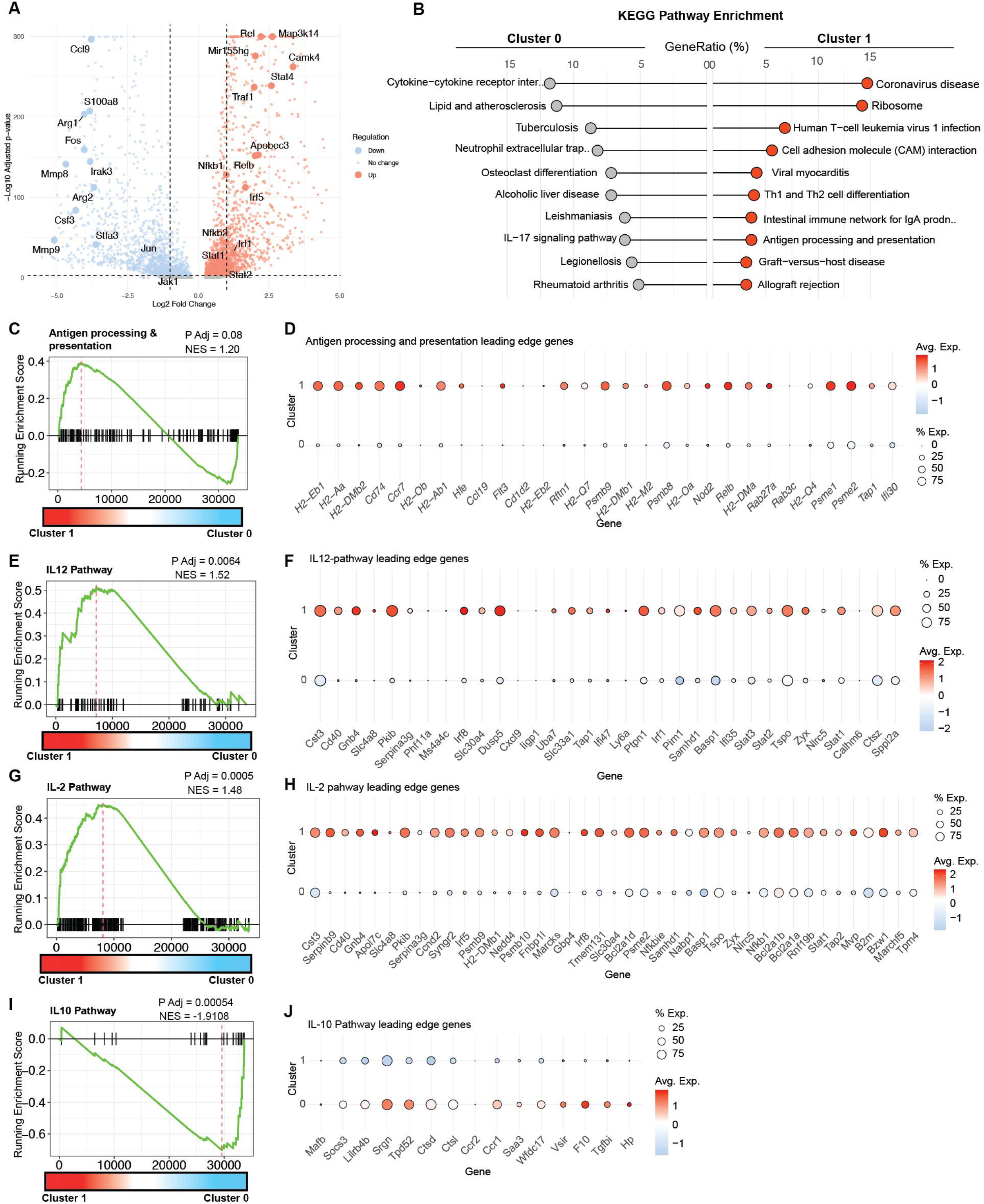
cDC1-like cluster 1 is enriched for type 1 immune polarizing gene programs. **A.** Volcano plot showing differential expression of signaling protein genes in cluster 1 vs cluster 0 identified from the scRNA-seq dataset. The X-axis represents log2 fold change and the Y-axis represents -log10 FDRq. Genes significantly enriched in cluster 1 are shown in orange, genes enriched in cluster 0 are shown in blue. Non-significant genes (FDRq>0.05) are indicated in gray. N=3 mice per scRNA-seq group. **B.** Lollipop plot of the top 10 pathways enriched in cluster 0 and cluster 1 based on KEGG pathway enrichment analysis of differentially expressed genes. **C**. Gene set enrichment analysis (GSEA) plot showing positive enrichment of antigen processing and presentation genes in cluster 1 compared to cluster 0. **D.** Dot plot showing the expression of leading-edge genes driving positive enrichment of antigen presentation pathways in cluster 1. Dot size indicates percentage of cells expressing the gene, color intensity represents scaled gene expression levels (red, highest gene expression and blue, lowest gene expression). **E.** GSEA plot showing positive enrichment of IL-12 pathway genes in cluster 1 compared to cluster 0. **F.** Dot plot showing expression of leading-edge genes driving the positive enrichment of the IL-12 pathway in cluster 1. **G.** GSEA plot showing the positive enrichment of IL-2 pathway genes in cluster 1 compared to cluster 0. **H.** Dot plot showing expression of leading-edge genes driving the positive enrichment of the IL-2 pathway in cluster 1. **I**. GSEA plot showing negative enrichment of the IL-10 pathway genes in cluster 1 compared to cluster 0. **J**. Dot plot showing expression of leading-edge genes driving negative enrichment of the IL-10 signaling pathway in cluster 1 compared to cluster 0.

### The mTORC1 signaling pathway is downregulated in cDC1-like cluster 1

We next dissected the molecular signaling pathways that establish this cDC1-like population by computing signaling pathway activity scores by cluster using PROGENy(*28*). Cluster 0 exhibited strong PI3K/mTOR and hypoxia signaling activity while JAK-STAT , WNT, p53, and VEGF signaling pathways were strongly activated in cluster 1 (**Figure 3A, S3A**). Because PI3K/mTOR signaling plays a critical role in governance of the IL-12 and IL-10 axis(*29, 30*), and exhibited strong differential signaling activity between cluster 0 and cluster 1, we examined this pathway in depth. Using the mTORC1 gene set derived from the Hallmark gene set collection(*31*), we performed GSEA, revealing that the mTORC1 pathway was significantly downregulated in cluster 1 compared to all other clusters (NES= **^-^**2.65) (**Figure 3B**). Further, the mTORC1 pathway was also significantly downregulated in cluster 1 in specific comparison to cluster 0 (NES = **^-^**1.22) (**Figure 3C**). This downregulation was driven by mTORC1 pathway genes that included *Ube2d3, Psmd12, Glrx, Ldha, P4ha1, Cd9, Pdk1, Egln3, Gsr, Slc711, Gbe1,* and *Itgb2* which were downregulated in cluster 1 compared to cluster 0 (**Figure 3D, S3B**). To examine whether the transcriptomic downregulation of the mTORC1 pathway translated to proteomic changes, we performed reverse phase protein array (RPPA) on BMDCs and measured global changes in protein level and phosphorylation status of over 300 signaling proteins. Supporting the inhibition of the mTORC1 pathway, BMDCs loaded with homologous MHC epitopes displayed a significant reduction in abundance and phosphorylation of mTORC1 pathway proteins including p-p70S6k, Ulk1, p-Raptor, Iκκb, p-Ulk1, c-Fos, NF-κBp65, PRAS40, & p-Alk (**Figure 3E, S3C**). To validate these BMDC findings in human moDCs, we loaded healthy donor-derived moDC with MHC epitopes derived from influenza hemagglutinin A which have been previously validated(*13, 14*). These consisted of homologous MHC peptides (MHC class I: **WLTGKNGKL** and MHC class II: RNLL**<u>WLTGKNGKL</u>**YPN) and control peptides made heterologous to each other through two G→M amino acid substitutions on the class II peptide (MHC class I: **WLTGKNGKL** and MHC class II: RNLL**<u>WLT</u>**M**<u>KN</u>**M**<u>KL</u>**YPN). After 24h maturation, phosphorylation of mTORC1 pathway proteins was ascertained by western blot. Consistent with mouse BMDC RPPA, we found significant reductions in phosphorylated mTOR (Ser2448) (**Figure 3F**) and its downstream targets p-p70S6K(Thr389)(*32*) (**Figure 3G**) and p-cFos(Ser32)(*30, 33*) (**Figure 3H**) among moDCs loaded with homologous MHC epitopes in comparison to control. Because the kinase activity of mTOR is dependent on its recruitment to the lysosomal membrane(*34*), we next examined the impact of homologous MHC epitope loading on lysosomal mTOR recruitment by measuring mTOR colocalization with the lysosome by confocal microscopy. Colocalization of mTOR with the lysosome was significantly reduced following moDC loading with homologous peptides in comparison to heterologous peptide control (**Figure 3I**). Further analysis using super resolution Stochastic Optical Reconstruction Microscopy (STORM) confirmed these findings at molecular (10-30 nm) resolution, with homologous MHC epitope loading of moDCs inducing the lengthiest nearest neighbor distance between mTOR and lysosomal signals (**Figure 3J**). Taken together, these transcriptional, proteomic, and spatiotemporal changes identified the mTORC1 pathway as a central player in establishment of the cDC1-like phenotype induced by loading of moDC with homologous MHC peptides.

**Figure 3.**
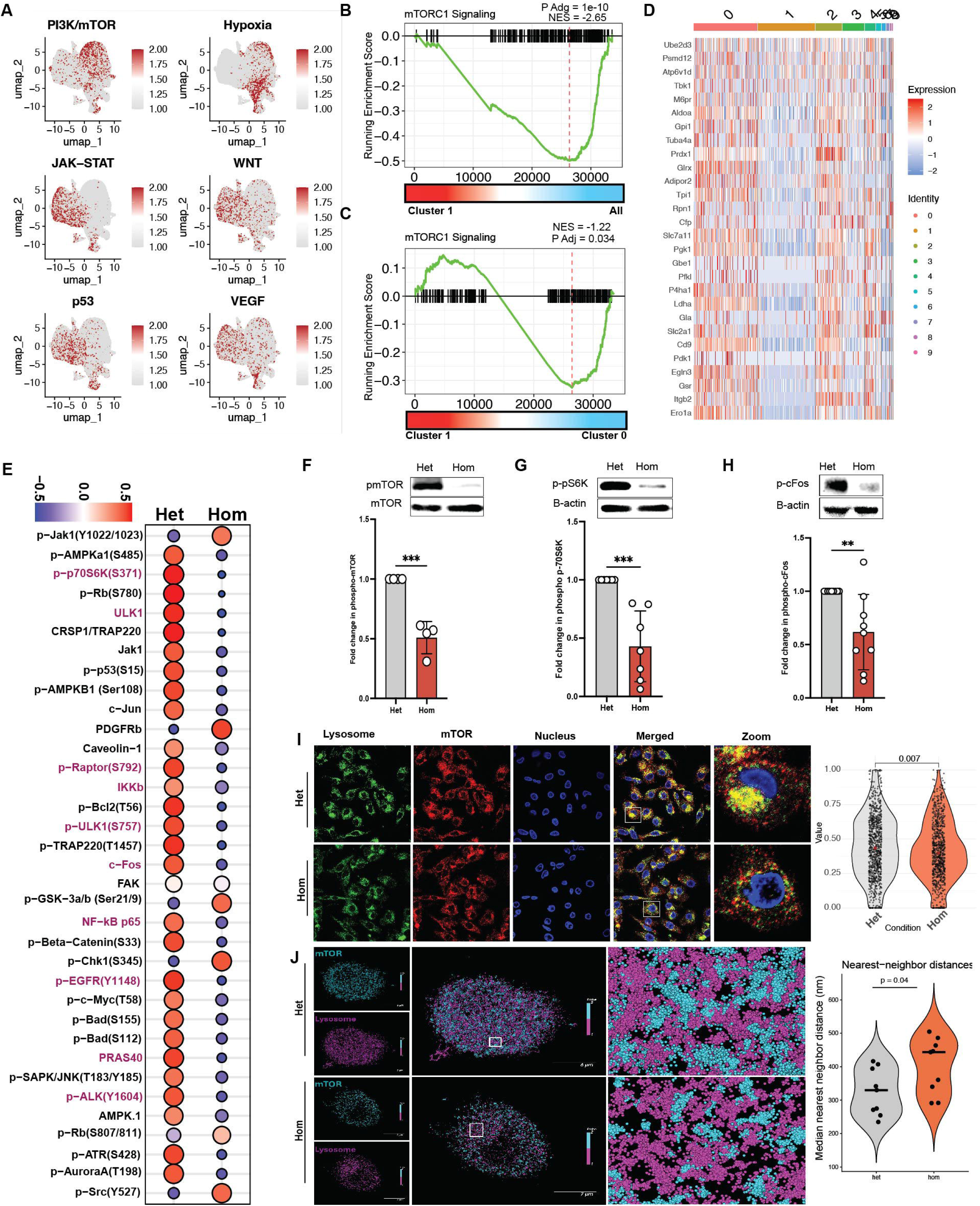
mTORC1 pathway signaling is downregulated in the cDC1-like cluster. **A.** UMAPs depicting signaling pathway activity inferred by PROGENy applied to the scRNA-seq dataset. Color indicates relative pathway activity score across cells (red, highest signaling activity; gray, lowest signaling activity). **B.** GSEA plot showing negative enrichment of mTORC1 signaling pathways genes in cluster 1 compared to all clusters. **C.** GSEA plot depicting negative enrichment of mTORC1 signaling pathway genes in cluster 1 compared to cluster 0. **D**. Heatmap showing expression of leading-edge genes contributing to the negative enrichment of the mTORC1 pathway in cluster 1. **E**. Dot plot of reverse phase protein array data showing levels of total and phosphorylated proteins in BMDC under indicated MHC peptide loading conditions. Proteins associated with mTORC1 pathway signaling are highlighted in pink. Data are a composite of N=4 mice. Statistical analysis performed by Student’s paired two-tailed t-test. **F**. Western blot analysis showing phosphorylated mTOR levels with corresponding densitometric quantification. N=4 healthy human donors. **G.** Western blot analysis showing phosphorylated p70S6K levels with corresponding densitometric quantification. N= 6 healthy human donors. **H.** Western blot analysis showing phosphorylated cFos levels with corresponding densitometric quantification. N=9 healthy human donors. Statistical analysis performed by Student’s paired two-tailed t-test. ***P<0.001, **P<0.01. **I**. Confocal microscopy images showing colocalization of mTOR with the lysosome (Lamp2) and corresponding quantification. N=5 healthy human donors. Statistical analysis performed by Wilcoxon rank-sum test. **J**. STORM super-resolution microscopy showing spatial proximity between mTOR and the lysosome (Lamp2) with quantification of nearest-neighbor distance between mTOR and lysosomal signals. Data are a composite of n=3 healthy human donors. Statistical analysis performed by Wilcoxon rank-sum test.

### Type 1 immune polarization is driven by NF-κB-mediated transcriptional reprogramming downstream of mTORC1 inhibition

A defining feature of cDC1 is an ability to produce high levels of IL-12 while keeping IL-10 expression low, supporting a role in driving type 1 immune responses. The mTORC1 pathway plays a central role in regulating the IL-12/IL-10 axis in DCs. Specifically, mTORC1 signaling promotes the expression and phosphorylation of c-Fos which enhances IL-10 gene transcription by directing NF-κB binding to the IL-10 promoter while reducing its binding to the IL-12a promoter(*29, 30, 35*). Since the mTORC1 pathway is significantly inhibited in the cDC1-like cluster, we hypothesized that moDC loading with homologous antigenic epitopes should inhibit the mTORC1 pathway, driving IL-12a gene expression via NF-κB (**Figure 4A**). To test this hypothesis, we performed ChIP-seq of NF-κB p65, the mTOR-regulated subunit that controls proinflammatory cytokine production in myeloid cells(*30, 35*). In homologous MHC epitope-loaded BMDCs, NF-κB was recruited to sites mostly located in the intergenic and intron regions, with significant differences in binding observed in the intron region of BMDC loaded with homologous MHC epitopes compared to the control (**Figure 4B**). In the regulation of proinflammatory gene expression, NF-κB cooperates with other transcription factors like Irf1 and Irf3 to recruit cofactors that govern gene transcription(*21, 36, 37*). To ascertain the relevance of this paradigm as a regulatory mechanism, we performed motif enrichment using HOMER(*38*). The most enriched binding motifs corresponded to proinflammatory transcription factors NFAT, IRF3, IRF8, NFAT:AP1 complex, ETV4, and NFATC2 (**Figure 4C, S4A/B**). To validate the regulatory role of these transcription factors in the cDC1-like cluster, we computed transcriptional activity scores using DoRothEA(*39*), an R package that uses curated transcription factors and their targets to infer transcriptional activity. The transcriptional activity of IRF8 and NFAT was highest in cluster 1 while the ETV4 and NFAT:AP1 complex (Fos and Jun) activity was highest in cluster 0 (**Figure 4D, S4C**). IRF3 and NFATC2 showed equivalent activity in both clusters 0 and 1 (**Figure S4C**). These findings demonstrate differential gene regulatory mechanisms engaged by NF-κB, depending on the homology of MHC peptides loaded on BMDCs. We next analyzed NF-κB binding to the IL-12a gene locus, noting a significantly enhanced peak in a cis regulatory element (CRE) located +7 kb upstream of the promoter (**Figure 4E**) in DC loaded with homologous epitopes. Concordantly, *IL-12a* gene expression and IL-12 secretion was consistently and significantly upregulated among homologous MHC epitope loaded DCs (**Figure 4F/G**). While *IL-12b* gene expression was increased (**Figure S4E**), no significant NF-κB binding was detected in the *IL-12b* gene locus (**Figure S4D**). In direct contrast, NF-κB binding to CREs located +4.6 kb upstream of the IL-10 promoter and 1 kb downstream the distal exon region was detected only among control DC loaded with heterologous peptide controls (**Figure 4H**). Consistently, *IL-10* gene expression and secretion were both significantly reduced among moDCs loaded with homologous peptide antigens (**Figure 4I/J**) in comparison to moDC loaded with heterologous control peptides. We next performed KEGG pathway enrichment analysis to identify signaling pathways differentially regulated by NF-κB. In keeping with our hypothesis, NF-κB regulated pathways related to type 1 immune responses in homologous MHC peptide loaded BMDCs (**Figure 4K**) compared to the control group (**Figure 4L**). The enrichment of viral infection pathways together with antigen presentation pathways in the homologous MHC peptide loaded group indicate a transcriptional program mediating the type 1 polarizing phenotype seen in moDCs after homologous epitope loading. Since antiviral responses correspond to gene programs that initiate cytotoxic T-cell responses, we identified the ChIP peaks driving the overrepresentation of the Human Papilloma Virus pathway and found significant binding peaks corresponding to critical genes that included *Itga3, Itgb7, Stat1, Creb1, Itga5, Dlg1, Gsk3b, Atp6v1g3, psmc1, Ifnar1, Grb2, Itgb8, Mapk1,* and *Itga1* among others (**Figure 4M**). Collectively, the data reveal a model in which inhibition of the mTORC1 pathway in response to homologous epitope loading drives NF-κB type 1 polarization of moDCs by enhancing *Il-12a* expression, inhibiting *Il-10* expression, and regulating other type 1 immune polarizing gene programs.

**Figure 4.**
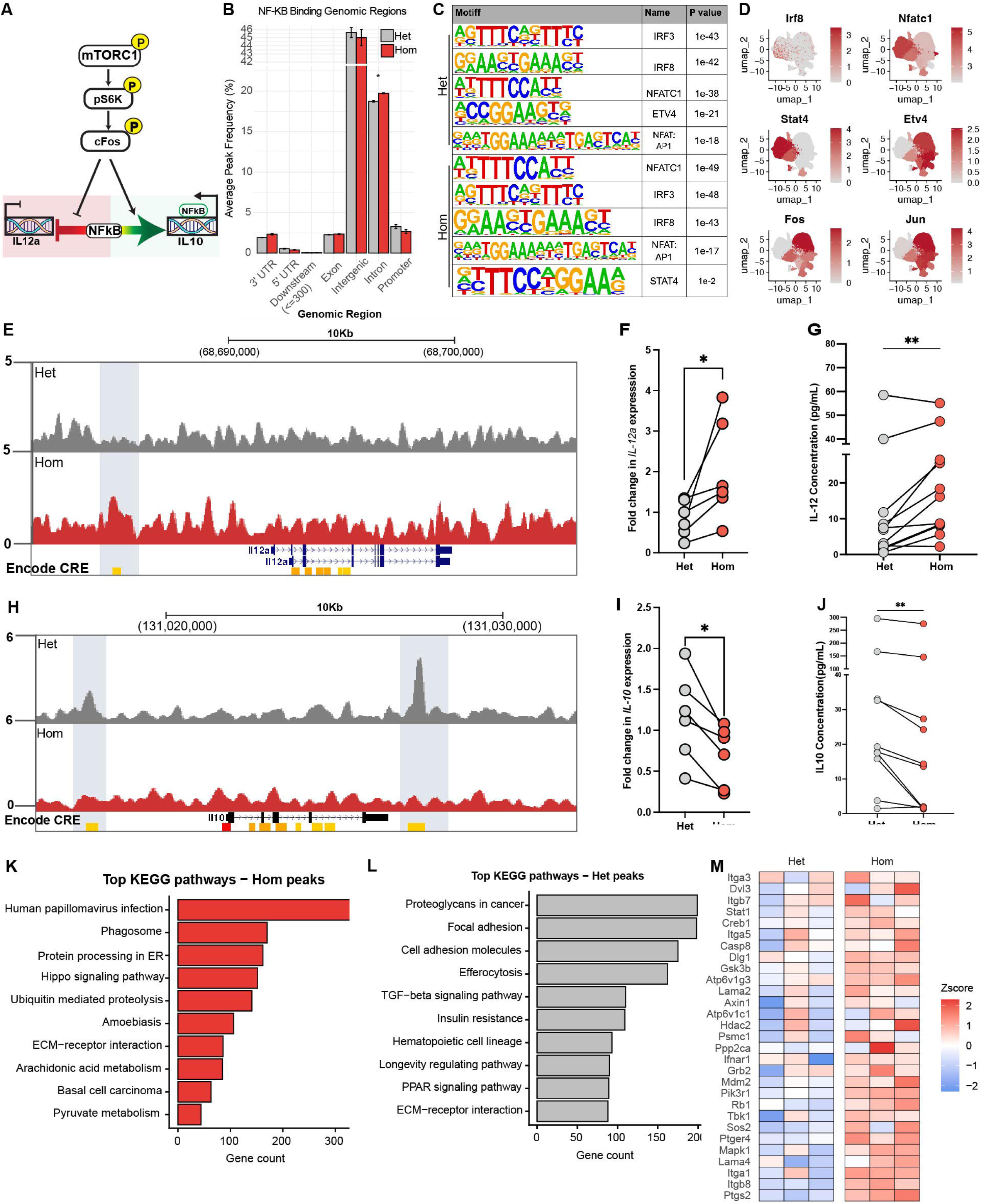
Type 1 immune polarization is driven by NF-κB-mediated transcriptional reprogramming downstream of mTORC1 inhibition. **A.** Conceptual model illustrating mTOR regulation of the IL-12/IL-10 axis in dendritic cells. Phosphorylated mTOR functions as a serine/threonine kinase that phosphorylates p70-S6K, which drives expression and subsequent phosphorylation of downstream target c-Fos. Phosphorylated c-Fos translocates to the nucleus where it inhibits NF-κB binding to the *Il12* promoter while inducing NF-κB binding to the IL-10 promoter, thereby driving *Il10* transcription. **B**. Bar plot showing differential NF-κB genomic binding locations in homologous epitope-loaded BMDCs compared with heterologous epitope loaded controls. **C.** Transcription factor binding motifs enriched within NF-κB peaks in the different peptide loading conditions as determined by Homer. **D.** UMAPs showing inferred activity of selected transcription factors identified by HOMER motif analysis, projected onto scRNA-seq clusters using DoRothEA-based transcription factor activity inference. **E.** Representative genome browser tracks (mm10 annotation) showing NF-κB ChIP-seq occupancy at *Il12a* locus. N=3 mice. **F.** IL-12a gene expression as determined by qPCR. N=6 healthy donor samples from healthy human donors pooled from 3 independent experiments. *P<0.05, **P<0.01 by Student’s paired two-tailed t -test. **G.** IL-12 protein secretion as measured by ELISA. N=10 samples from healthy human donors pooled from 3 independent experiments. *P<0.05, **P<0.01 by Student’s paired two-tailed t test. **H.** Representative genome browser tracks (mm10 annotation) showing NF-κB ChIP-seq occupancy at the *Il10* locus. N=3 mice. **I**. IL-10 gene expression as determined by qPCR. N= 6 samples from healthy human donors pooled from 3 independent experiments. *P<0.05, **P<0.01 by Student’s paired two-tailed t test. **J.** IL-10 protein secretion as measured by ELISA. N=10 samples from healthy human donors pooled from 3 independent experiments. *P<0.05, **P<0.01 by Student’s paired two-tailed t test. **K-L.** KEGG pathway enrichment analysis of genes associated with NF-κB binding sites identified in homologous MHC peptide loaded BMDC (K) and heterologous MHC peptide loaded DC (**L**). Pathways are ranked by adjusted p value and gene count. **M**. Split heatmap showing leading-edge genes driving the enrichment of the human papillomavirus infection pathway in homologous antigen loaded BMDC. B-C, H, K-M. BMDCs differentiated from C57BL/6 mice were loaded with homologous or heterologous MHC peptides and cultured for 24 hours. ChIP was performed using NF-κBp65 antibody and coprecipitated DNA was sequenced. F-G, I-J. monocytes from healthy human donors were differentiated into moDCs and loaded with homologous or heterologous MHC peptides. After 24-hour maturation, ELISA and qPCR were performed.

### Inhibition of mTOR enhances the cDC1-like phenotype and drives moDC-mediated type 1 immune responses in T-cells

To confirm the role of the mTORC1 pathway in establishment of the cDC1-like phenotype, we inhibited mTOR in BMDCs loaded with homologous or heterologous MHC peptides and quantified the expression of cDC phenotypic markers by flow cytometry and cytokine secretion by ELISA (**Figure 5A**). We first noted that mTOR inhibition with rapamycin does not affect DC maturation as demonstrated by similar levels of CD80 (**Figure S5A**) and only a modest increase in CD86 (**Figure S5B**) expression in BMDCs treated with rapamycin. We next assessed expression of Xcr1 and CD11b as markers for cDC1-like and cDC2-like cells, respectively. We determined that Xcr1 expression significantly increased among BMDCs treated with rapamycin irrespective of MHC peptide homology (**Figure 5B/C**). Conversely, CD11b was significantly reduced in both groups when mTOR was inhibited (**Figure 5D/E**), suggesting biased differentiation of BMDCs toward a cDC1-like state when mTOR is inhibited. Because mTOR regulates the IL-12/IL-10 axis in DCs, we wondered whether direct mTOR inhibition might skew this axis toward IL-12 expression. We found that IL-12 secretion significantly increased with rapamycin treatment across all groups (**Figure 5F**). Likewise, IL-10 expression was significantly reduced across the treatment groups following mTOR inhibition (**Figure 5G**). Increased expression of Xcr1 and IL-12 suggested an enhancement of the cDC1-like phenotype in BMDCs when the mTORC1 pathway is inhibited by rapamycin. To determine whether mTOR-inhibited moDCs drive type 1 biased T-cell differentiation, we co-cultured allogeneic PBMCs with rapamycin-treated human moDCs and measured IFN-γ expression as well as T-cell subset markers by flow cytometry (**Figure 5H**). IFN-γ production was significantly increased in PBMCs co-cultured with rapamycin-treated moDC (**Figure 5I/J**) and further, there was a significant increase in Tbet^+^ CD4^+^ (**Figure 5K/L**) and Granzyme+ CD8^+^ T-cells (**Figure 5M/N**) with a concomitant increase in Tbet^+^ CD8^+^ T-cells (**Figure S5C**). Additionally, we observed significant reductions in PD-1 expression in both CD8^+^ (**Figure S5D**) and CD4^+^ T-cells (**Figure S5E**) as well as GATA3 expression among CD4^+^ T-cells (**Figure S5F**). These findings suggest a type 1 biased differentiation among T-cells cocultured with rapamycin-treated moDC and collectively confirm that inhibition of mTOR plays a critical role in establishment of the cDC1-like phenotype in moDCs, enhancing their ability to drive type 1 immune responses.

**Figure 5.**
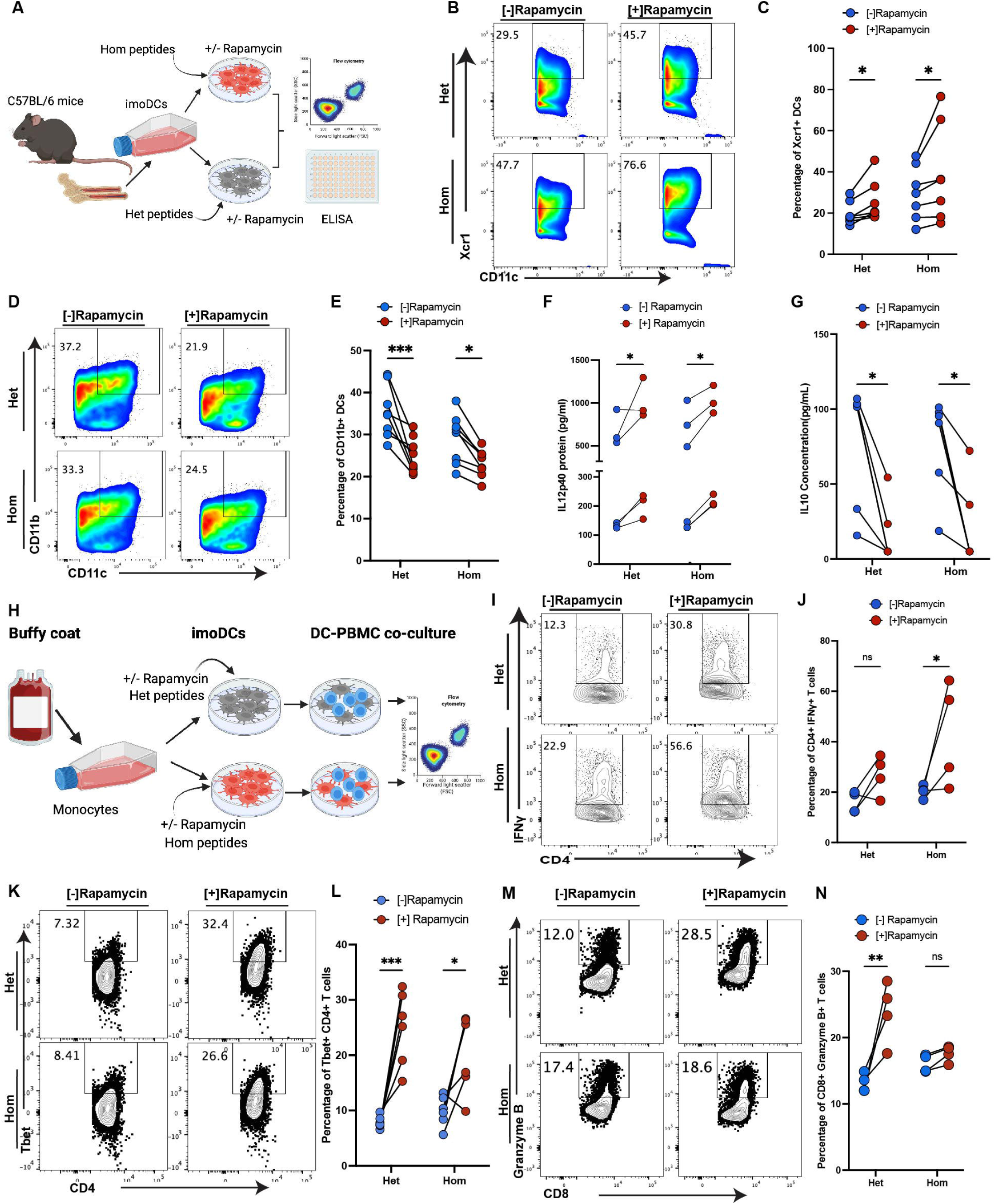
Inhibition of mTOR enhances the cDC1-like phenotype and drives DC-mediated type 1 immune responses in T cells. **A.** Experimental workflow for ex vivo inhibition of mTORC1 signaling in BMDCs. BMDCs were differentiated from C57BL6 bone marrow and loaded with either homologous or heterologous LCMV MHC binding peptides and mTORC1 signaling was inhibited with rapamycin. Cells were subsequently analyzed by flow cytometry, and cell culture supernatants were collected for IL-12 and IL-10 quantification. **B.** Representative FACS plots displaying Xcr1 expression in BMDC loaded with MHC peptides with or without rapamycin treatment. **C.** Quantification of Xcr1 expression. N= 7 mice pooled from 2 independent experiments. **D.** Representative FACS plots displaying CD11b expression in BMDC loaded with MHC peptides with or without mTOR inhibition. **E**. Quantification of CD11b expression +/- mTOR inhibition. N=8 mice pooled from 2 independent experiments. **F**. IL-12 secretion in supernatants of BMDC loaded with MHC peptides +/- mTOR inhibition. N=6 mice pooled from 2 independent experiments. **G**. IL-10 expression in BMDC supernatants +/- mTOR inhibition. N=6 mice pooled from 2 independent experiments. **H**. Experimental workflow for *ex vivo* stimulation of PBMCs with MHC peptide-loaded moDCs +/- mTOR inhibition. Immature moDC derived from healthy donors were loaded with MHC peptides and treated with rapamycin to inhibit mTOR signaling. DC were co-cultured with allogeneic PBMCs from healthy donors. After 14 days, T-cell subsets were analyzed by flow cytometry. **I**. Representative flow cytometry plots showing IFNγ^+^ CD4^+^ T-cells in DC-PBMC co-cultures. **J.** Quantification of IFNγ^+^ CD4^+^ T cells. N=4 healthy donors. **K**. Representative flow cytometry plots showing Tbet^+^ CD4^+^ T-cells in DC-PBMC co-cultures. **L**. Quantification of Tbet^+^ CD4^+^ T-cells. N=5 healthy human donors. **M.** Representative flow cytometry plots displaying Granzyme B^+^ CD8^+^ T-cells in DC-PBMC co-cultures. N=4 healthy donors. p < 0.05, ∗∗p < 0.01, and ∗∗∗p < 0.001 by two-way ANOVA with post-hoc analysis by Sidak’s multiple comparison test.

### The cluster 1 cDC1-like gene signature is enriched among human clinical trial moDC vaccine products

To evaluate the potential translational relevance of cDC1-like moDC differentiation, we analyzed bulk RNA-seq data from human clinical moDC vaccine products, loaded in a manner that enforces homologous antigenic loading(*40–42*), that had been administered to phase I clinical trial patients being treated for glioblastoma (NCT04552886) and pancreatic ductal adenocarcinoma (NCT04157127) (**Figure 6A**). Read counts were compared between the clinical vaccine product and autologous, patient-specific control moDC loaded in heterologous fashion. In comparison to autologous control DC loaded in heterologous fashion, clinical vaccine products exhibited a gene profile consistent with the cDC1-like functional phenotype, upregulating significantly *BATF3, XCL2, PSMB3, PSMC5, PSME2, INF2, SERPINB6, IL32, LAD1, NDUFA13*, and *CLEC18A*, while downregulating anti-inflammatory genes such as *THBS1, TGFBR2, MERTK, ACVRIC* and *IRF2* (**Figure 6B**). To determine whether homologous antigenic loading induces a cDC1-like transcriptional profile similar to that identified in BMDCs loaded with homologous LCMV peptides, we conducted fast gene-set enrichment analysis (FGSEA) using the cDC1-like cluster gene set comprised of the top 1,000 genes expressed in the cDC1-like cluster. Consistent with mouse BMDC scRNA-seq data, the cDC1-like cluster gene signature was significantly enriched in human moDCs loaded with homologous antigens in comparison to control with a net enrichment score (NES) of 1.69 and a p value of 0.00037 (**Figure 6C**). To examine the transcriptional changes at the individual patient level, we calculated a paired cDC1-like gene set enrichment scores for each sample. The cDC1-like gene signature was enriched in 88% (29/33) of products (**Figure 6D**), suggesting a translational relevance. The leading edge genes driving positive enrichment included genes related to cDC1 identity and antigen presentation (*Batf3, Psme1, Psmb7, Psmb8, Psme2, Psmc5, Cd74, Uba7, Uba52 and Cst3*), inflammatory signaling (*Selplg, Nfkbie, Ifi35, Il20rb, Aif1, and Il2rg*) cytoskeletal remodeling and migration (*Ccl17, Pfn1, Myl6, Myo1g, Pdlim4, S100A4, S100A10, Lad1, Lst1, Tesc* and *Arfp1*) and genes related to the cellular stress response, immune cell priming, and homeostasis (*Serpinb6b, Pla1a, Lad1 and Nsmce1)* as first characterized in the mouse BMDC scRNA-seq data (**Figure 6E**), and recapitulated among human clinical moDC vaccine products (**Figure 6F, S6**). Because mTORC1 pathway inhibition was critical for establishment of the cDC1-like phenotype seen in mouse BMDCs, we examined the mTORC1 pathway enrichment by FGSEA, noting significant downregulation of the mTORC1 pathway with a net enrichment score of -1.59 among DC loaded with homologous antigens (**Figure 6G**). This downregulation was observed in 94% (31/33) of moDC vaccine products (**Figure 6H**), cementing the central role of mTORC1 pathway inhibition in establishment of the cDC1-like phenotype. Collectively, these findings reveal a previously unappreciated mechanism through which moDC are reprogrammed towards a cDC1-like fate.

**Figure 6.**
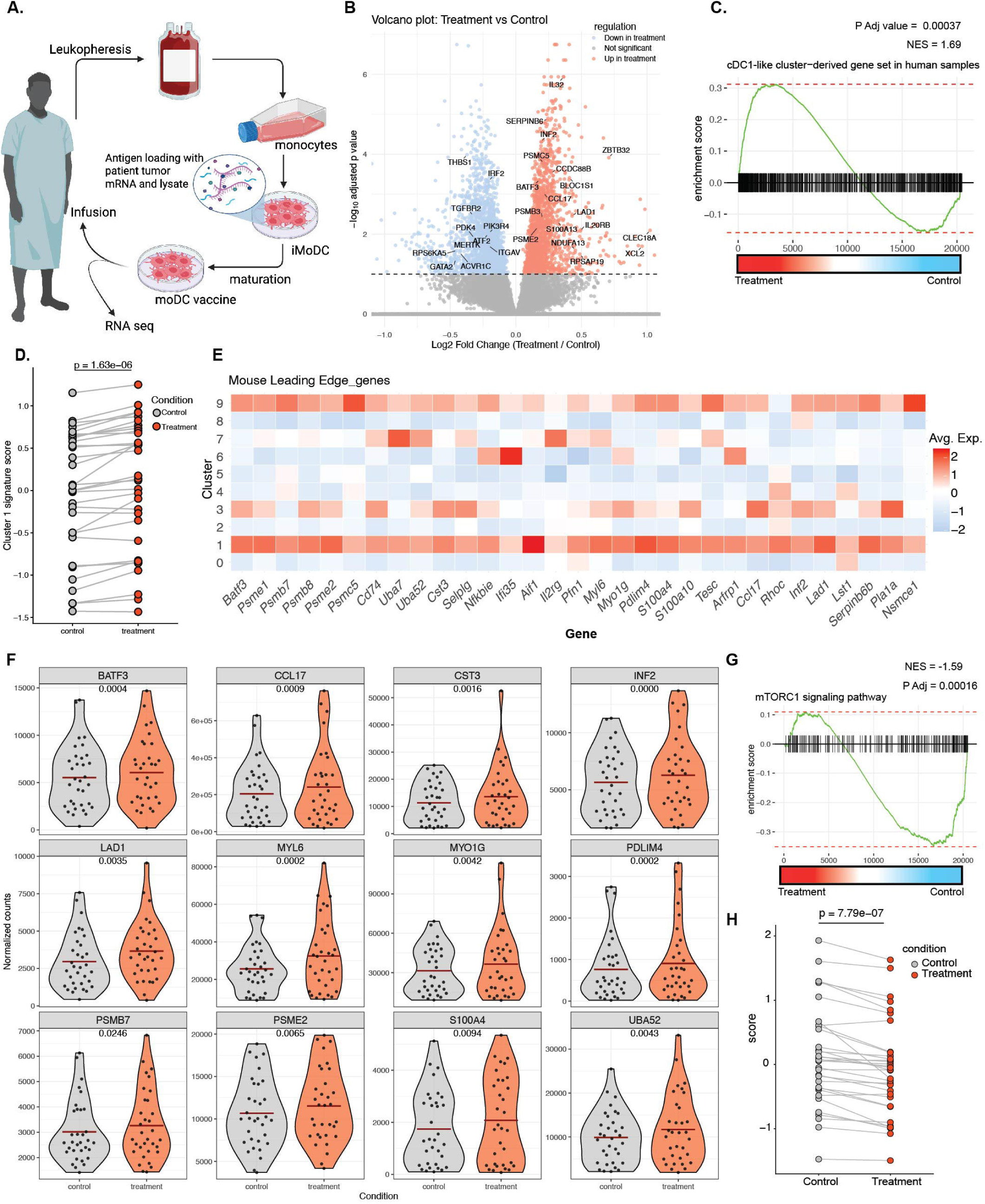
The cluster 1 cDC1-like gene signature is enriched among human clinical trial moDC vaccine products. **A.** Schematic outlining experimental design for moDC homologous antigen vaccine preparation for treatment of glioblastoma and pancreatic ductal adenocarcinoma patients. In total, 33 patient vaccines were included in the bulk RNA-seq data analysis. **B.** Volcano plot showing differentially regulated genes between each homologous antigen-loaded moDC vaccine and patient-specific autologous control loaded in heterologous fashion. **C.** GSEA plot showing positive enrichment of the mouse cDC1-like cluster-derived genes among human moDC vaccine products. **D.** Paired dot plot showing the cDC1-like gene enrichment score in homologous antigen-loaded moDC vaccine products compared to the heterologous antigen-loaded control. Statistical analysis was performed by Student’s paired two tailed t-test. **E.** Heatmap showing the expression of leading-edge in BMDC scRNA-seq data. **F.** Violin plots of select leading edge genes driving positive enrichment of the cDC1-like cluster gene set in human moDC vaccines. Statistical analysis performed by Student’s paired two-tailed t-test. **G.** GSEA plot depicting negative enrichment of mTORC1 signaling pathway genes in human moDC vaccine products. **H.** Paired dot plot showing the mTORC1 signaling pathway gene score in homologous antigen-loaded moDC vaccine products compared to the heterologous antigen-loaded control. Statistical analysis performed by Student’s paired two-tailed t-test.

## Discussion

Viral infection generates homologous class I and class II MHC epitopes that strongly activate CD4 and CD8 T cell responses(*7, 8, 10*), providing a potential blueprint for therapeutic manipulation of moDC. We have shown that these homologous MHC epitopes are not a mere byproduct of viral infection, but a true signal – a pattern that can be recognized to induce a type 1 polarizing phenotype in moDCs(*13, 14*). Indeed, moDC vaccines based on this strategy have been shown to induce T-cell memory responses in glioblastoma patients(*43–45*). However, the mechanistic basis of these empirical observations has yet to be characterized. Here, we show that the loading of moDC with homologous MHC epitopes induces differentiation of a cDC1-like phenotype characterized by upregulated canonical cDC1 markers including *Batf3, Irf8, Zeb1, Zfp366,* and *Xcr1* along with upregulated viral infection response pathways. Importantly, the data show that establishment of this cDC1-like phenotype is mediated by inhibition of the mTORC1 pathway, facilitating expression of IL-12a and other type 1 polarizing genes via differential binding of NF-κB to inflammatory rather than tolerogenic chromosomal loci. Indeed, inhibition of mTOR using rapamycin enhanced the cDC1-like phenotype and boosted type 1 biased differentiation of T-cells when rapamycin-treated moDCs were co-cultured with allogeneic PBMCs. Lastly, this work reveals that the cDC1-like transcriptional program is conserved from mouse to man and can be enhanced in human clinical moDC vaccine products generated by means of a homologous antigen-based loading strategy(*43–45*).

cDC1s are the specialized subset of dendritic cells known to initiate type 1 immune responses(*22, 46*). Their development is mediated by a coordinated network of transcription factors including Batf3, Irf8 and Id2(*47–49*). More recent work has additionally identified the transcription factors Zfp366 and Zeb1 as critical to the regulation of cDC1 development and homeostasis(*23, 50*). Consistent with these developmental and regulatory programs, the induced cDC1-like population identified in the present study exhibited high expression of *Batf3, Irf8, Zfp366* and *Zeb1*. Additionally, *Ly75* (DEC-205), a C-type lectin endocytic receptor critical for antigen presentation and reported to be highly expressed on CD8a^+^ cDC1(*4, 51–53*), was also highly expressed in the cDC1-like cluster 1. While *Xcr1* transcripts were minimal in scRNA-seq data, flow cytometry showed high expression of Xcr1 in homologous MHC epitope loaded BMDC compared to control. Furthermore, KEGG pathway enrichment revealed high expression of viral infection pathways in the cDC1-like cluster, suggesting a type 1 immune polarizing transcriptional program consistent with the type 1 immune responses orchestrated by cDC1. Indeed, this cluster exhibited high expression of genes related to antigen presentation pathways, IL-12 and IL-2 signaling pathways, and a significant reduction in the expression of genes related to IL-10 and IL6-Jak-Stat3 signaling pathways, exemplifying the classical type 1 immune polarizing response characteristic of the cDC1 subset(*4*).

The DC type 1 immune polarizing phenotype is tightly regulated by a coordinated network of signaling pathways to ensure appropriateness of response. The mTORC1 pathway is one of the key pathways critical to identity and phenotype in part through regulation the IL-10/IL-12 axis in antigen presenting cells(*30, 35, 54*). When mTORC1 is active, p-cFos displaces NF-κB from the IL-12 promoter, redirecting its binding specificity to the IL-10 promoter and fostering a tolerogenic phenotype(*29, 30, 35*). However, inhibition of mTOR skews the axis toward IL-12 gene expression, thereby favoring type 1 immune polarization(*30, 35, 55*). Here, we demonstrate that the mTORC1 pathway is significantly downregulated in the identified cDC1-like cluster compared to all other clusters of BMDC differentiating in response to homologous or heterologous antigenic loading. Moreso, this inhibition was further evident at the proteomic level as demonstrated by a decrease in expression and phosphorylation of mTORC1 pathway related proteins including p-p70S6K, ULK1, p-raptor, Ikkb, p-ULK1, c-FOS, PRAS40, & p-ALK. The recruitment of mTOR to the lysosome was also significantly reduced in moDCs loaded with homologous MHC epitopes. As mTOR recruitment is required for its kinase activation by lysosomal membrane resident small G protein Rheb(*34, 56, 57*), inhibition of lysosome recruitment may substantially contribute to inhibition of mTOR phosphorylation and initiation of downstream inhibitory effects. Consistent with this hypothesis, ChIP-seq indicated a shift in NF-κB binding dynamics, with binding to a CRE +7 Kb upstream of the IL-12a promoter in homologous MHC loaded DCs and differential binding to CREs located +4.6 kb upstream of the IL-10 promoter and 1 kb from the distal exon region of IL-10 gene among DC loaded in heterologous fashion. These results align with previous studies reporting that inhibition of mTOR enhances IL-12 gene expression via NF-κB (*30, 35*) and highlights a complex regulatory network governing the DC IL-10/IL-12 axis.

In clinical practice, rapamycin is used as an immunosuppressant to prevent allograft rejection, most prominently through its impact upon Treg expansion (*58, 59*). However, strong experimental evidence supports an additional, proinflammatory role for rapamycin in antigen-presenting cells(*30, 35, 60*), observations supported by the present work. Inhibition of mTOR using rapamycin enhanced the cDC1 phenotype characterized by an increase in Xcr1 expression and IL-12 secretion and concomitant reduction in CD11b expression and IL-10 secretion. When co-cultured with T-cells, these rapamycin-treated moDCs enhanced the differentiation of IFNγ^+^CD4^+^ T-cells, Tbet^+^CD4^+^ T-cells, Tbet^+^CD8^+^ T-cells, and granzyme B^+^CD8^+^ T-cells, suggesting enhanced ability to induce cytotoxic T-cell responses. This is consistent with previous reports that inhibition of mTOR with rapamycin enhances IL-12 expression in dendritic cells and enhances type 1 immune responses in vivo (*35, 60*). These findings may provide mechanistic rationale for incorporation of *ex vivo* mTOR inhibition to enhance the efficacy of moDC-based vaccines.

Overall, our work reveals a previously undescribed mechanism of immune governance in which homologous MHC epitopes serve as a type 1 polarizing signal that induces the differentiation of a cDC1-like population from moDCs. The development of the cDC1-like population is dependent on mTORC1 pathway inhibition, a strategy that can be leveraged to enhance the efficacy of moDC-based therapies.

## Acknowledgments

We thank Dr. Toby Lawrence and Dr. Ammar S. Cheema for generously sharing their R script for pseudotime analysis and members of our laboratory for the insightful scientific discussions and advice. This work was supported in part by National Institutes of Health grants NIH R01 AI127387 and NIH R01 AI153326 (to WKD), and BRASS (Baylor Research Advocates for Student Scientists) grant (to SBA). This study was further supported in part by the Baylor College of Medicine flow cytometry core facility under the academic leadership of Dr. Christine Beeton and the expert assistance of Mr. Joel M. Sederstrom. This study was additionally supported in part by the Genomic and RNA Profiling Core at Baylor College of Medicine with funding from NIH S10 grant 1S10OD023469, NCI grant P30CA125123, CPRIT grant RP200504. We thank the Epigenomics Profiling Core (EpiCore) at MD Anderson Cancer Center for help with ChIP assays. AKJ is partially supported by institutional support to EpiCore. Lastly, this study also received support from the Single Cell Genomics Core at Baylor College of Medicine with funding from CPRIT RP250580, NIH S10OD032189 and P30CA125123.

## Author contributions

WKD and SBA conceived the project and designed experiments. SBA conducted experiments, performed omics data analysis and image analysis. AM, KJE, NB, ZNB, JVP, WL, AT, OSD, AP, and LU performed DC-antigen loading, gene expression, flow cytometry, and western blot experiments. MS and VA designed the protocol for STORM imaging. VA performed STORM image acquisition. SH, SZ, and MHN performed reverse phase array sample processing and data analysis. MJ conducted single cell library preparation. KP and DCK performed sequencing. AKJ and PA performed ChIP and ChIP library preparation. VK designed experiments and performed DC antigen loading experiments. WKD designed and supervised the studies and provided funding. SBA and WKD co-wrote the paper with input from VK. All authors read and approved the final version of the manuscript.

## Data and materials availability

Sequencing data has been uploaded to the Gene Expression Omnibus (GEO) database with accession numbers GSE325592 (scRNA-seq), GSE324817 (ChIP-seq), and GSE325601 (bulk RNA seq). All other data needed to evaluate the conclusions of this work will be made available upon reasonable request to the corresponding author.

## Supplementary Materials

Materials and methods

Figures S1 to S6

Table S1

---

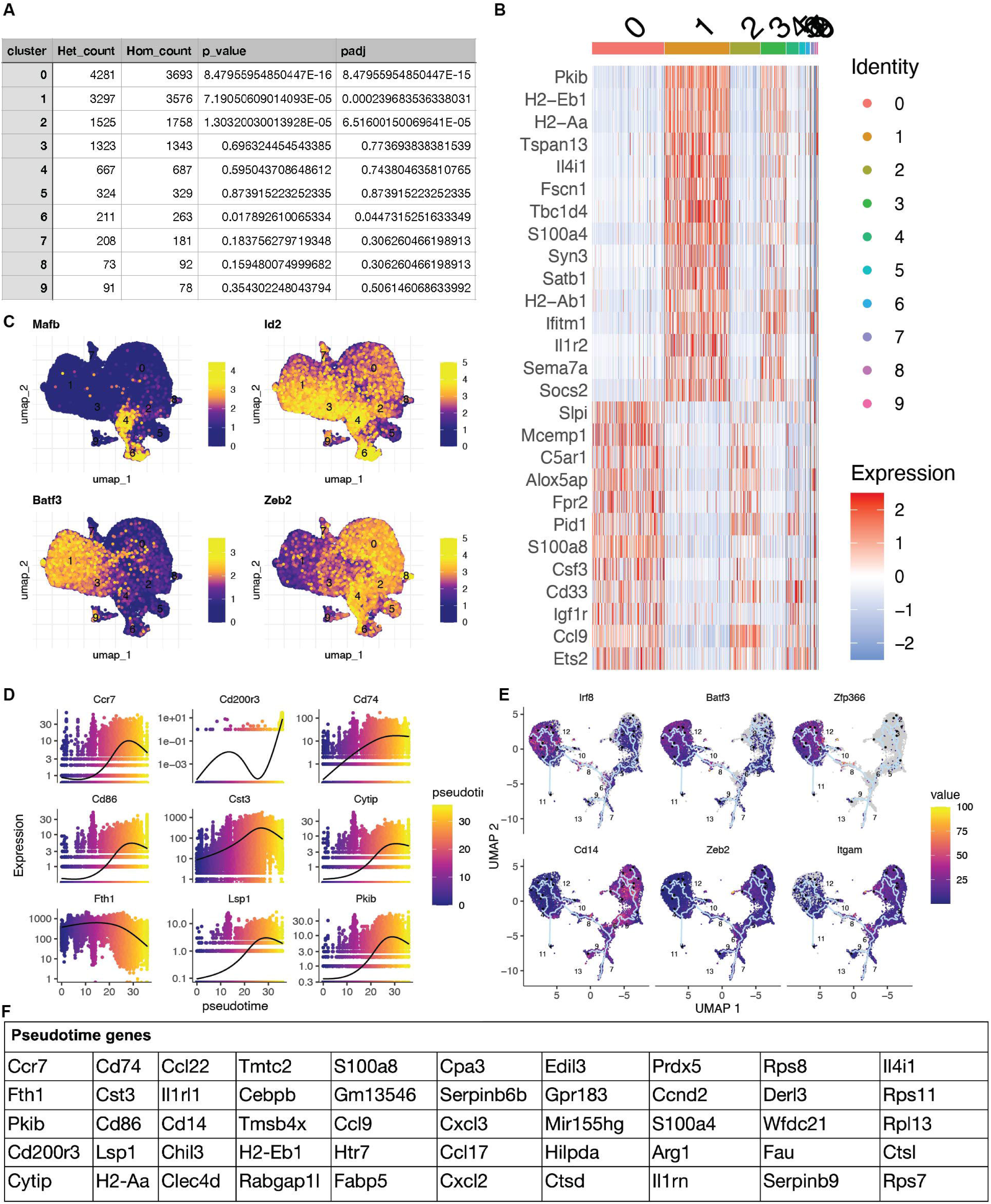

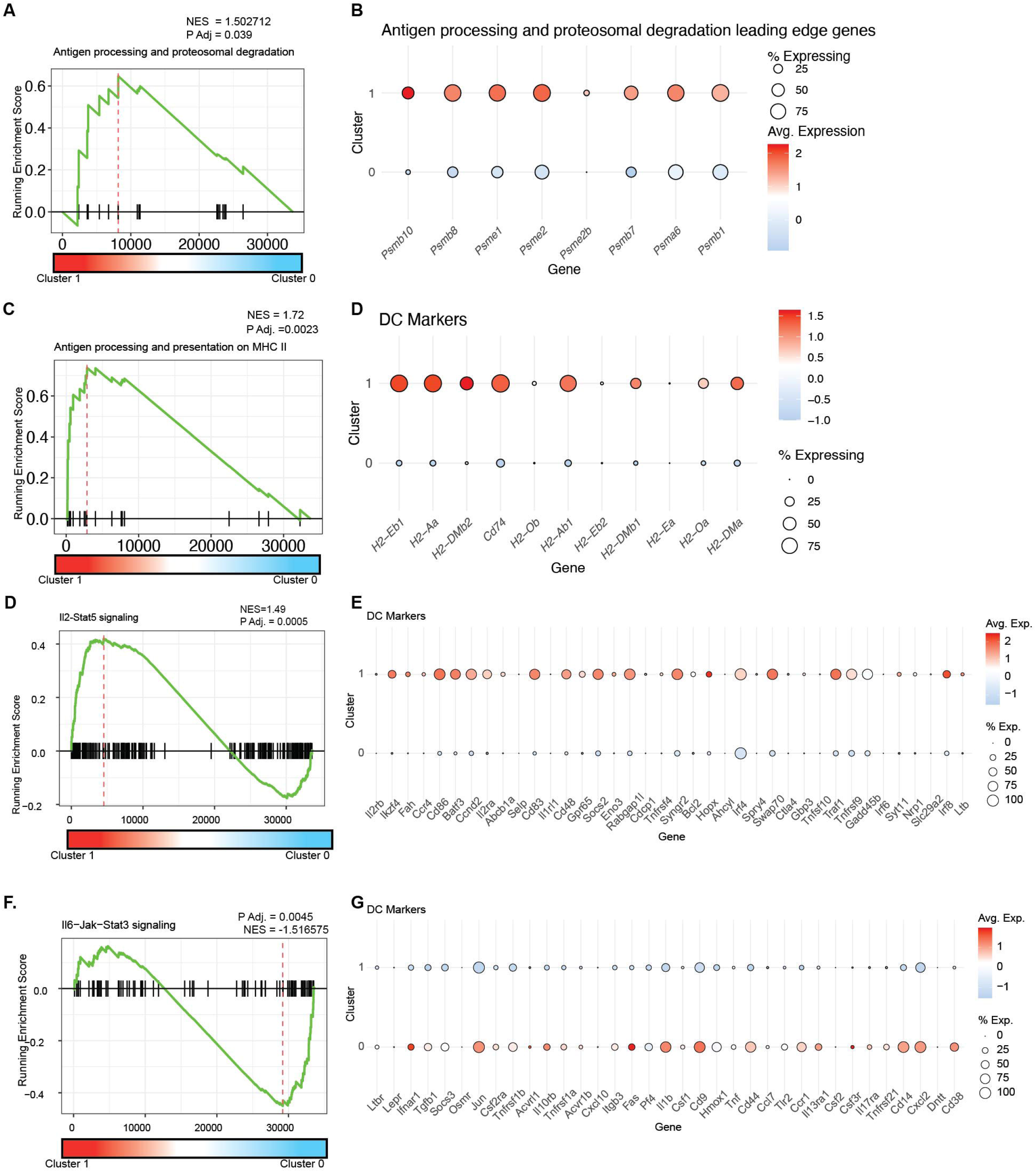

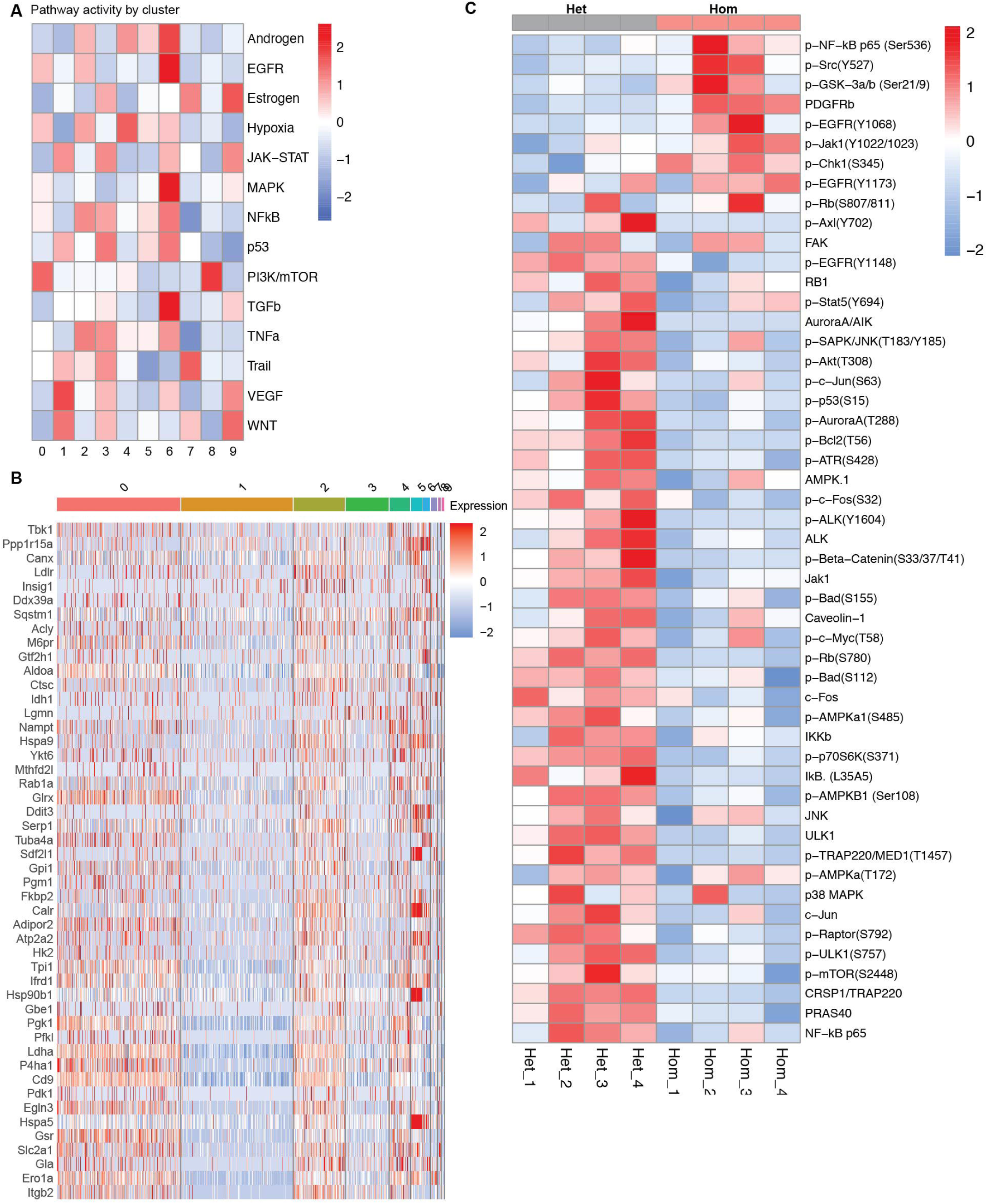

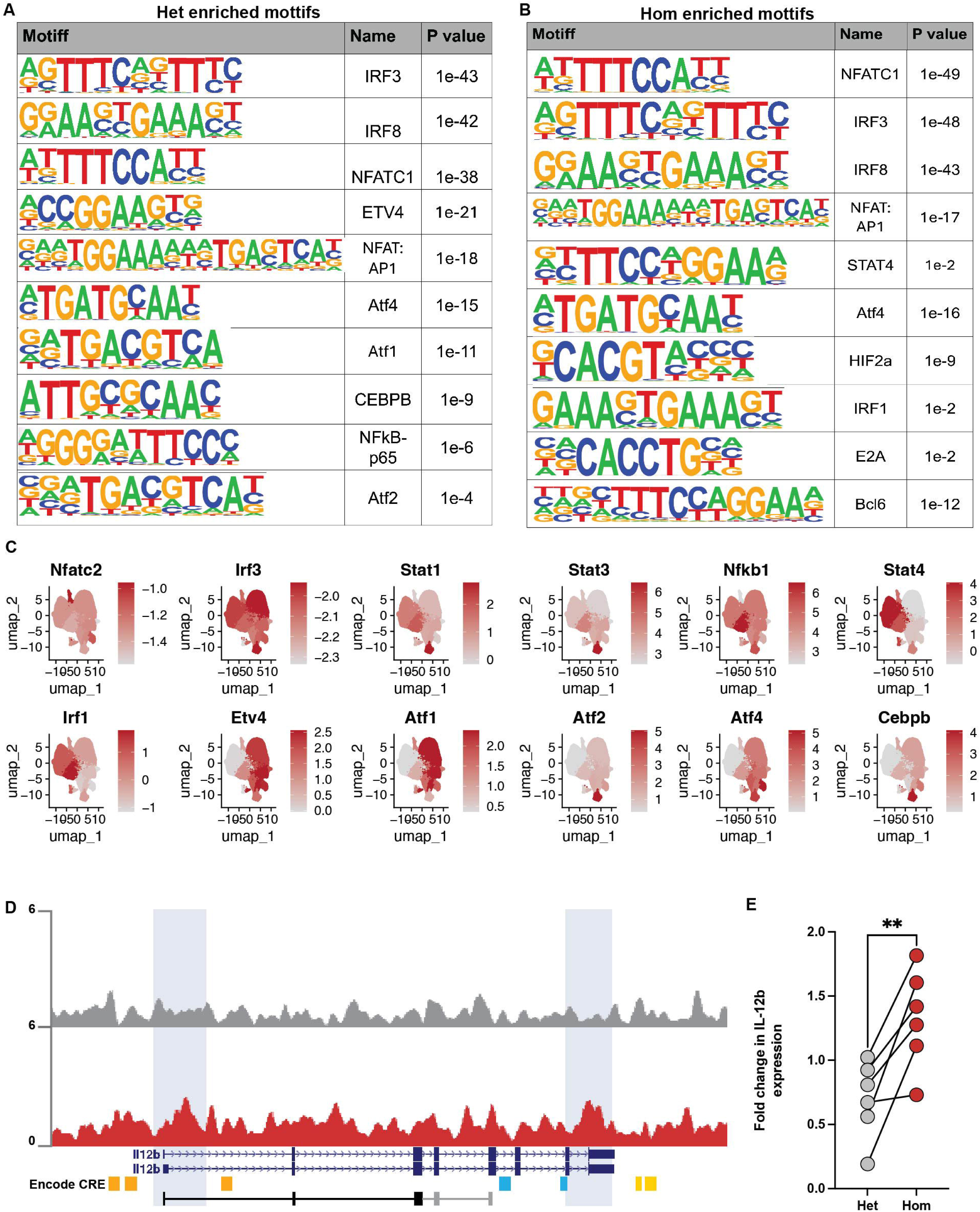

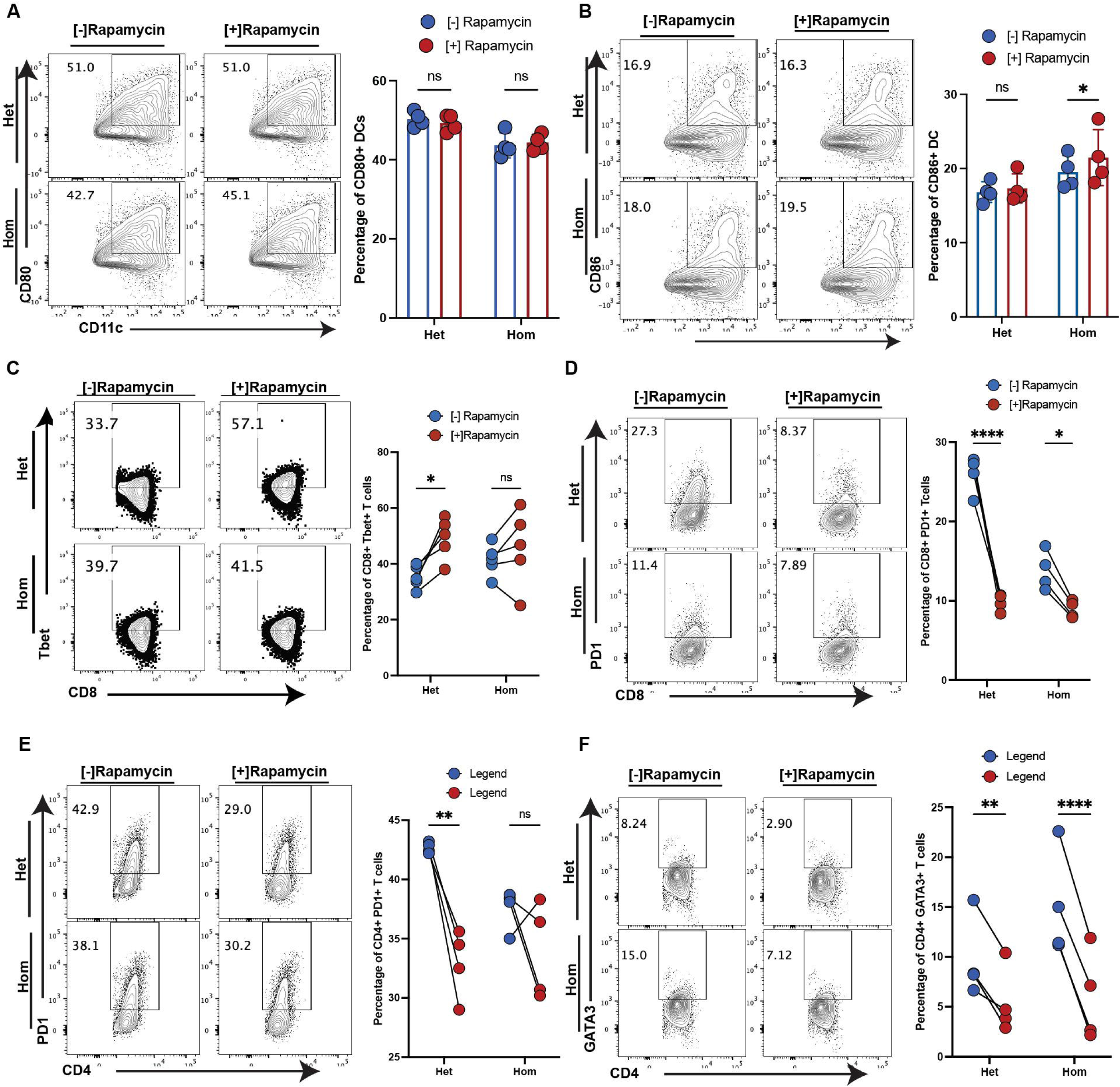

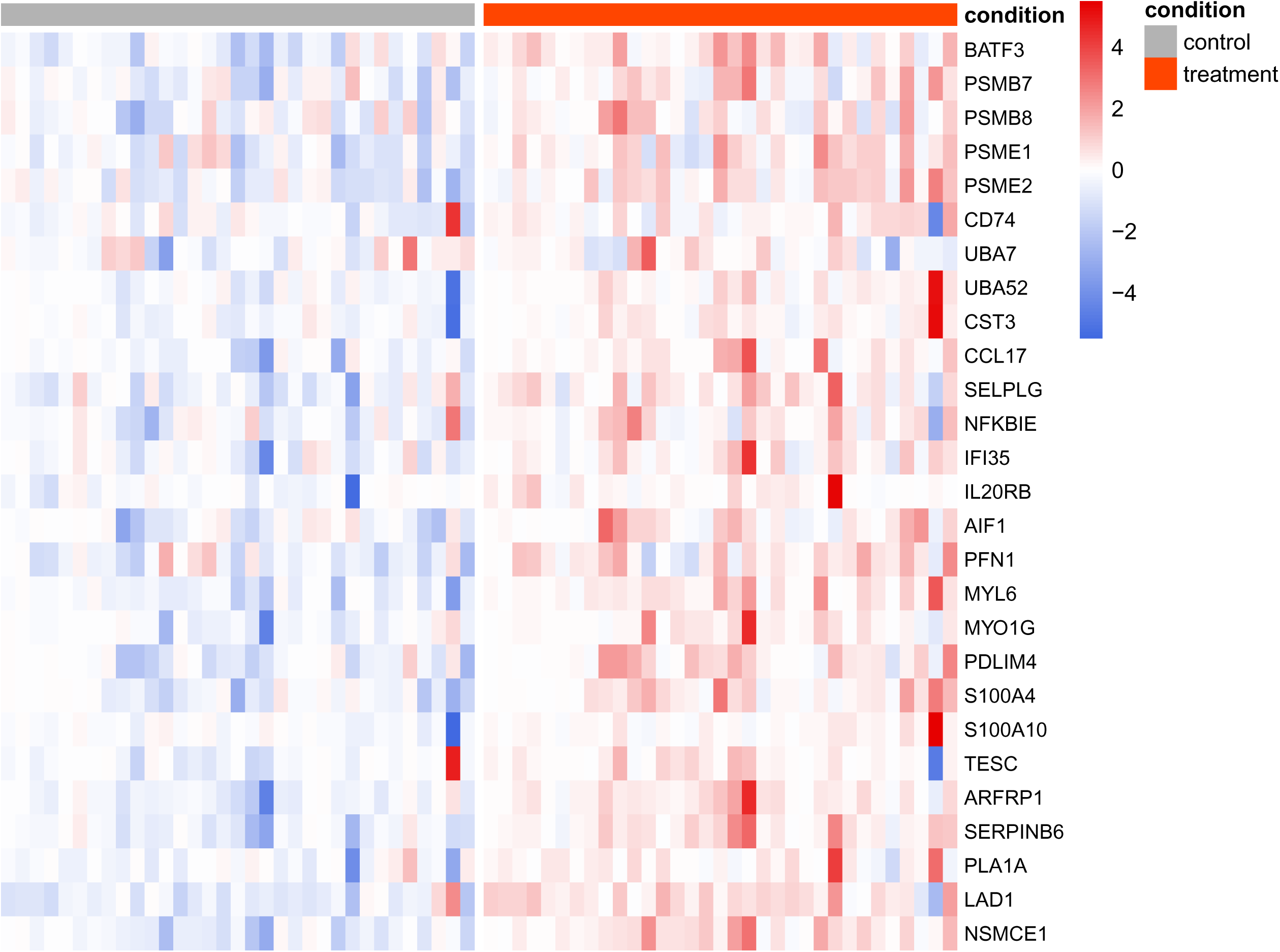

