## Supplementary material for "Overlapping MHC class I/II Epitopes Program cDC1-like Differentiation of Monocyte-Derived Dendritic Cells via mTORC1 Signaling Inhibition": Materials and methods, and supplimentary figure legends

**Methods**

**Mouse BMDC Isolation**. C57BL/6 mice at 6-8 weeks of age were euthanized and dissected to obtain femurs and tibias. Long bones were placed bone marrow-side down in a porous 0.5 ml tube inside a 1.5 ml Eppendorf tube and centrifuged at 5000 xg for 15 minutes. Contaminating red blood cells were lysed with ACK buffer (Gibco, #A1049201) for 10 minutes, filtered through a 70μm filter, and washed twice with PBS. Marrow cells were counted and plated at 2 x 10^6^/ml in AIM V medium (Gibco, #12055083) supplemented with 10% mouse serum (Sigma-Aldrich, #M5905) and 1% antibiotic-antimycotic (Gibco, #15240062), 30 ng/ml GM-CSF (R&D systems, # 6130-GR-050), and 10 ng/ml IL-4 (R&D systems, # 404-ML-025). BMDCs were differentiated for 6 days, with media and cytokine replenishment on days 3 and 5.

**Human dendritic cell differentiation**.

Buffy coats were obtained from healthy donor blood donations at the Gulf Coast Blood Center (Houston, Texas) in accordance with approved protocols. Peripheral blood mononuclear cells (PBMCs) were isolated by density gradient centrifugation in Lympholyte cell separation medium (Cedarlane, #CL5010) from buffy coat diluted 1:4 in PBS. CD14^+^ monocytes were isolated by immunomagnetic isolation with CD14 microbeads according to the manufacturer’s instructions (Miltenyi, #130-050-201). DC differentiation was induced by addition of 50 ng/ml GM-CSF (R&D Systems, #215-GM) and 10 ng/ml IL-4 (R&D systems, #BT-004-050) to monocyte cultures seeded at 2x10^6^ cells/ml in AIM V supplemented with 10% human serum (Valley Biomedical Products and Services, #HP1022HI) and 1% anti-anti (Gibco, #15240062). Media and cytokines were replenished on day 3 and day 5. Immature DCs were harvested on day 6 of culture using cell dissociation buffer (Gibco, #13151014).

**Peptide loading and maturation**. Immature DCs were loaded with MHC peptides as previously characterized and described(*1*). LCMV glycoprotein peptides: gp33–41 (KAVYNFATC), gp31–45 (GIKAVNFATCGIFA), and gp66–80 (DIYKGVYQFKSVEFD), and Influenza A New Caledonia hemagglutinin peptides: WLTGKNGL, RNLLWLTGKNGKLYPN, and RNLLWLTMKNMKLYPN, all synthesized by United Bio Systems (Herndon, VA).

Briefly, Immature DCs were washed with PBS and resuspended in SERVATOR B Solution (Benchchem, #B611682) at 40 x 10^6^ cells/ml. 500 μl of the cell solution was transferred to pre-chilled electroporation cuvettes (Biorad, #1652088). Thereafter, MHC peptides were added at a concentration of 5μg/ml and vortexed vigorously for 30sec. The cuvettes were incubated on ice for 10 minutes and thereafter electroporated using a Gene-Pulser Xcell System (Biorad, #1652660) set at exponential waveform with 300 V, 150 μF, ∞ resistance, and 4 mm gapped cuvette. Electroporated cells were immediately transferred to AIM V supplemented with 2.5% serum and resuspended at a concentration of 3 x 10^6^/ml. After a 3 hour incubation at 37^º^ C, cells were supplemented with AIMV containing 7.5% serum, 1% anti-anti, and DC maturation cocktail consisting of: human (IL-4; 10 ng/μl, GM-CSF; 50 ng/μl, IL-1β; 10 ng/μl, TNF; 20 ng/μl, IL-6; 15 ng/μl, and PGE_2_; 1 μg/μl), and mouse (IL-4; 10 ng/μl, GM-CSF; 30 ng/μl, IL-1β; 10 ng/μl, TNF; 20 ng/μl, IL-6; 15 ng/μl, and PGE_2_; 1 μg/μl). The DCs were then incubated at 37^º^ C until harvest at 24 or 48 hours.

**mTOR inhibition.** Rapamycin (Sellekchem, #S1039) was added at a concentration of 25 nM at the time maturation reagents were added. After a 6-hour incubation, DCs were washed to remove rapamycin, and resuspended in complete medium supplemented with maturation cytokines. The DCs were then incubated at 37^º^ C until harvest at 24 or 48 hours.

**RNA Isolation and real-time quantitative PCR.** Following the addition of maturation cocktail, DCs were incubated at 37º C in a humidified chamber supplemented with 5% atmospheric CO_2_ and harvested at 24 hours for IL-12a and IL-10 gene expression assays. RNA was extracted using TRIZOL^TM^ (Invitrogen, #15596018) according to the manufacturer’s instructions. For cDNA synthesis, 1 μg RNA was used to synthesize cDNA using the High-Capacity cDNA Reverse Transcription kit (Applied Biosciences, #43-688-14) according to manufacturer’s instructions. The PCR conditions for cDNA synthesis were as follows: 10 min incubation at 25° C, 120 minute incubation at 37°C, and a 5-minute termination step at 85° C. For gene expression, 20 ng cDNA was used for qPCR amplification using the 7500 Real-Time PCR System (Applied Biosystems). Each sample was analyzed in triplicate using the 18S rRNA transcript to normalize expression levels and a gene-specific probe listed in Table S1. Relative gene expression was calculated using the 2^-ΔΔCT^ method and was presented as fold change to control.

**RNA-seq data analysis.** We analyzed RNA-seq data generated from analysis of human clinical moDC vaccine products for patients enrolled in phase I clinical trials for glioblastoma (NCT04552886) and pancreatic ductal adenocarcinoma (NCT04157127) (*2-4*). Differential gene expression analysis was performed using the DESeq2 package in R. Gene set enrichment analysis (GSEA) was performed using clusterProfiller package in R to determine enrichment of the mouse cDC1-like cluster gene signature and the mTORC1 pathway.

**ChIP-seq.** After 24h maturation, DCs were harvested and washed thoroughly with PBS. Cells were cross-linked using 16% ChIP-grade formaldehyde (Thermo Fisher Scientific, #28908) and incubated at room temperature for 12 minutes. The cross-linking reaction was stopped with 2.5M glycine at room temperature for 5min. Samples were then washed three times with ice-cold PBS, and chromatin immunoprecipitation (ChIP) assays were performed at the MD Anderson Cancer Center Epigenomics Profiling Core following a protocol previously described(*5, 6*) with certain modifications. Chromatin was prepared by nuclear isolation followed by lysis. Lysates were subjected to sonication with a Bioruptor Pico (Diagenode) to obtain DNA fragments ranging from 200-600 bp followed by centrifugation at 16,000 xg for 10 minutes at 4° C. The supernatant containing chromatin lysate was pre-cleared with Dynabeads^TM^ Protein A beads (Invitrogen, # 10-001-D) for 1 hr at 4° C and then incubated with NF-κB P65 antibody (Cell Signaling Technologies, #8242) conjugated with Dynabeads^TM^ Protein A beads overnight at 4° C. The following day, beads were collected using DynaMag-2 magnet (Invitrogen, # 12-321-D), washed extensively, and ChIP DNA was isolated after reversing the crosslinks. Input and ChIP DNA libraries were prepared using NEBNext® Ultra™ II DNA Library Prep Kit for Illumina® (New England Biolabs, Ipswich, MA) following the manufacturer’s instructions. ChIP-seq libraries were pooled in equimolar fashion and 150 pmol were loaded onto a single lane of a NovaSeq 6000 S4 flow cell (Illumina, p/n 20028317), based on qPCR quantification using a ViiA7™ Real-Time PCR System. The library was prepared following the XP Workflow protocol (Illumina kit p/n 20021664) and amplified by exclusion amplification onto a nanowell-designed, patterned flowcell using the Illumina NovaSeq 6000 sequencing platform. The PhiX Control v3 adapter-ligated library (Illumina p/n FC-110-3001) was spiked in at 2% by weight to ensure balanced diversity and to monitor clustering and sequencing performance. A paired-end 150 bp cycle run was performed to sequence the flow cell on the NovaSeq 6000 sequencer. An average of 156 million read pairs per sample was sequenced.

**ChIPseq data analysis.** FASTQ sequences were subjected to quality control using FastQC. This was followed by alignment to the mouse reference genome mm10 using bowtie2(*7*). The aligned reads were processed for sorting and duplicate removal using SAMtools(*8*) and Sambamba(*9*). The deduplicated reads were then normalized by random downsampling to equalize sequencing depth across samples and indexed using SAMtools. Bigwig files were generated using deeptools (*10*) and visualized on the UCSC Genome Browser(*11*). Significant ChIP-seq signal enrichment peaks were identified using the Model-based Analysis of ChIP-Seq (MACS2)(*12*). ChIPSeeker(*13*) and ClusterProfiler(*14*) were used for peak annotation and pathways enrichment analysis, while the enriched transcription factor binding motifs were identified using Homer(*15*).

**Single cell RNA sequencing.** Following harvest, live/dead sorting was performed on a BD FACSAria (BD Biosciences), and cells were submitted to the Single Cell Genomics core at the Baylor College of Medicine for library preparation. Viable single cells were captured in droplet emulsions using the 10x Genomics Chromium X, targeting 5,000 cells per sample. scRNA-seq libraries were generated using the 10x Genomics Single-Cell 3′ Gene Expression on-chip multiplexing chemistry according to the manufacturer’s protocol (including barcoding, reverse transcription, cDNA amplification, and library construction) using the Chromium Single Cell 3’ Library Construction Kit (10x Genomics). Equimolar (100 pmol) 3′GEX v4 single-cell RNA-seq libraries were pooled and loaded onto a single lane of a NovaSeq X Plus 10B flow cell (Illumina, p/n 20085594) with 5% PhiX Control v3 spike-in. Libraries were sequenced on a NovaSeq X system using a paired-end 150 bp configuration, generating an average of 500 million read pairs per sample. FastQ file generation was executed using bcl2fastq, and QC reports were generated using CellRanger v5.0.1. FASTQ files were processed using the CellRanger count pipeline (CellRanger v10.0.0, 10X Genomics) and aligned to the mm10 mouse reference genome (*16*). Seurat version 4.4.1(*17*) was used to analyze the aggregated data. To identify high quality cells for downstream analysis, quality control was performed to filter out cells with mitochondrial genes >7.5%, less than 500 genes per cell, or greater than 9,500 genes per cell. The counts were normalized using the LogNormalize function in Seurat. We performed principal component analysis (PCA) to reduce the dimensionality of the data. The top 12 principal components were selected to construct the UMAP embeddings. Using the FindClusters function at a resolution of 0.2, we found 10 transcriptionally distinct clusters. The FindMarkers function was used to identify specific markers in the clusters. Pathway analysis was performed using enrichKEGG function in ClusterProfiler(*18*).

**Transcription factor activity inference.** To infer activity for HOMER-identified transcription factors in the single-cell RNA sequencing data, we used DoRothEA regulon resource and the decoupleR framework(*19, 20*). High confidence mouse TF-target interactions (confidence levels A-C) were obtained from the DoRothEA database. Gene expression data were filtered to retain only genes overlapping with the selected regulons. TF activities were inferred from pseudobulk average expression profiles using the univariate linear model (ULM) method implemented in decoupleR, with a minimum regulon size of five target genes. The resulting TF activity scores and associated p-values were adjusted for multiple testing by means of the Benjamini-Hochberg procedure. TFs of interest as identified by HOMER in the ChIP-seq data were selected for downstream analysis and visualization. TF activity was mapped back to individual cells based on cluster assignment on UMAP embeddings using Seurat.

**PROGENy signaling pathway activity inference.** We used PROGENy (1.28.0)(*21*) to infer signaling pathway activity across the single cell RNAseq clusters. The PROGENy framework was applied to the pseudobulk expression matrix with scaling enabled and the top parameter set to 300 pathway-responsive genes per pathway. The Signaling pathway activity was mapped back to individual cells based on cluster assignment on UMAP embeddings using Seurat.

**Pseudotime analysis.** Monocle (version 1.4.26)(*22*) was used as previously described(*23*) to investigate differentiation trajectories and dynamics of gene expression trajectories of moDC. Raw RNA count matrices were extracted from Seurat RNA assay and corresponding cell-level metadata and gene annotations were used to construct a monocle3 cell_data_set object. Cells were preprocessed using principal components analysis (PCA) and embedded into UMAP space. Clustering was performed on the UMAP embedding and the principal graph was learned to construct activation trajectories. Pseudotime was calculated by ordering cells along the principal graph from cluster 6, a biologically informed root cluster representing the earliest differentiation state. Pseudotime-associated gene expression changes were used to characterize dynamic transcription programs across clusters.

**Western Blot.** After 24 hour maturation, DCs were harvested and washed twice with PBS. Cells were then lysed in 200 μl of NP-40 lysis buffer [150 mM NaCl, 50 mM Tris-HCl, pH 7-8, 1% NP-40] and incubated on ice for 30 minutes. Thereafter, lysates were sonicated for 15 seconds at 40% power using the Q125 Sonicator (Qsonica, #Q125-110). Lysates were clarified at top speed for 15 minutes at 4^°^ C. Protein concentration in the supernatant was determined by Lowry assay using the RC DC Protein assay kit (Biorad, #5000121). Equal amounts of protein were loaded on SDS-polyacrylamide gel and transferred onto a 0.2 μm PVDF membrane (Biorad, #1620177). After confirming protein transfer by ponceau S staining (Thermo Scientific, #A40000279), membranes were blocked for 1 hour at room temperature in 5% PhosphoBlocker (Cell Biolabs, #AKR-104) buffer for phosphorylated proteins and 5% dry non-fat milk (Biorad, #1706404XTU) for non-phosphorylated proteins. After blocking, membranes were incubated with the appropriate primary antibody with shaking overnight at 4^°^ C. Membranes were washed 3 times in TBS-T and then incubated with the appropriate HRP-conjugated secondary antibody with shaking at room temperature for 1 hour. Membranes were washed 3 times in TBS-T and visualized in Clarity Western ECL Substrate (Biorad, #1705061). Proteins with a weak signal were visualized using SuperSignal™ West Femto Maximum Sensitivity Substrate (Thermo Fisher Scientific, #34094). Images were acquired on a ChemiDoc Imaging System (Biorad, #170-8280) and quantified using Image lab software version 6.1.

**Reverse Phase Protein Array.** RPPA assays were performed as described(*24, 25*). Specifically, protein lysates from moDC samples were prepared using modified T-PER™ Tissue Protein Extraction Reagent (Thermo Scientific, #78510) supplemented with protease and phosphatase inhibitors (Thermo Scientific, #78440), diluted to 0.5 mg/mL, and denatured. Samples were arrayed in triplicate onto nitrocellulose membrane–coated slides (Grace Bio-Labs, #305177) using a Quanterix 2470 Arrayer (Quanterix) and probed on an Autolink 48 stainer (Agilent/Dako) with 300 antibodies targeting total and phosphorylated proteins. Detection used IRDye® 680RD streptavidin (LI-COR Biosciences, #926-68079), and total protein was assessed by SYPRO® Ruby Protein Blot Stain (Cat#S11791, Thermo Scientific). Slides were scanned using a GenePix 4400AL (Molecular Devices), background-subtracted, and normalized to total protein. Antibodies not meeting quality criteria were repeated or excluded as described(*26*) with modification. Normalized data was used for downstream analysis. Significantly altered proteins were identified by paired comparisons performed using two-tailed paired t-tests implemented in R. Resulting p values were corrected for multiple comparisons using the Benjamini-Hochberg false discovery rate (FDR) adjustment.

**ELISA.** ELISA was performed to measure the concentrations of IL-10, IL-12p70, and IL-12p40 in cell culture supernatant. Following 24-hour maturation, supernatants were harvested by centrifugation at 5,000xg for 5min. The supernatants were stored at ^-^80^o^C until the day of analysis. The human uncoated ELISA kit (Invitrogen, #88-7126-22 and cat# 88-7106-22,) and mouse uncoated ELISA kits (Invitrogen, cat# 88-7105-22 and cat# 88-7120-88) were utilized according to manufacturer’s instructions. Briefly, high binding 96 well plates (Corning, #9018) were coated overnight with capture antibody at 4° C. The next day, plates were washed three times with PBS containing 0.05% Tween 20 and blocked overnight in 200μl of kit-provided ELISA diluent according to the manufacturer’s instructions. Following blocking, the plate was washed once and 100 μl standard dilution and samples were added in duplicate. Plates were incubated at 4° C overnight and washed five times in PBS-T prior to incubation with biotin-conjugated detection antibody. Plates were incubated for 1 hour at room temperature, washed five times in PBS-T, and incubated for 30 minutes in streptavidin-HRP solution. Signal was developed using tetramethylbenzidine substrate at room temperature for 15 minutes and stopped by adding 100 μl stop solution. Absorbance signal at 450 nm was immediately acquired on a fluorescent microplate reader (Thermo Fisher Scientific).

**STORM Microscopy.** Cover slips ((#1.5H Thickness, Ø25 mm, ThorLabs) were UV sterilized and coated with poly-D-lysine hydrobromide (100 ug/uL, Sigma Aldrich, #P6407-5MG) and seeded with 1.5 x 10^6^ DC in complete medium in a 6-well plate (Corning, # 3516). After maturation, plates were centrifuged at 400 xg for 5 minutes, washed in PBS, and fixed with 4% paraformaldehyde (Thermo Fisher Scientific, # 043368.9M) in PBS. Slides were incubated in a blocking solution (5% normal donkey serum, 0.3% Triton X-100 in PBS) for 1 hour. Slides were then incubated using primary antibodies diluted in blocking solution overnight at 4° C (mTOR, Cell signaling technologies, #2983S and Lamp2, Abcam #ab25631). Coverslips were washed three times in PBS and incubated with fluorochrome conjugated secondary antibodies for 1 hour at room temperature. After washing, cells were stored in PBS at 4^°^ C until imaging. STORM imaging was performed on a Bruker Vutara 352 instrument (Bruker) using a 60X water objective (UPLSAPO60XW). Images were acquired for 5 cycles at 200 nm axial step size and 250 frames per step. Stained cover slips were mounted in a collared well with imaging buffer added on top of the sample. STORM imaging buffer composition used for image acquisition was made immediately prior to imaging and consisted of 20 μl 1 M MEA (Sigma), 10 μl 2-mercaptoethanol (Sigma), 20 μl 50x Gloxy in 950 μl 50 mM Tris-HCl (pH 8.0) + 10 mM NaCl + 10% (w/v) glucose. 50x Gloxy stock contained 8440 AU Glucose oxidase (Sigma) and 70200 AU Catalase (Sigma) and was prepared in 50 mM Tris-HCl (pH 8.0) + 10 mM NaCl. Raw localization data from each image was clustered using the Ordering Points to Identify the Clustering Structure (OPTICS) algorithm on Vutara software. All clustered images were analyzed using a general particle distance of 0.2 μm and a particle count per cluster of 20 for both channels. Nearest neighbor distances were computed using a custom python script based on pandas and scikit-learn. Location coordinates were filtered to remove low confidence detection and thereafter filtered datasets were reduced to three-dimensional spatial coordinates (x,y,z) and corresponding localization estimates. Nearest-neighbor analysis was performed using NearestNeighbors implementation from scikit-learn module(*27*). For each localization, the distance to the nearest neighboring location was computed in three-dimensional space using Euclidean distance metrics. Differences in median nearest-neighbor distance between DC loaded with homologous MHC epitopes and heterologous peptide control were assessed using Wilcoxon rank-sum test.

**Confocal Microscopy.** 24-well glass bottom plates (15.0mm, 1.5H, Cellvis, #P24-1.5H-N) were coated with poly-D-lysine hydrobromide (100 μg/μl, Sigma Aldrich, #P6407-5MG) and seeded with 1 x 10^6^ DC. After 24-hour maturation, cells were centrifuged and then fixed in 4% paraformaldehyde in PBS for 15 minutes. After washing three times in PBS, he cells were incubated in blocking solution (5% normal goat serum, 0.3% Triton X-100 in PBS) for 1 hour at room temperature. Primary antibodies (mTOR Cat#2983S and Lamp2 Cat#ab25631) diluted in blocking solution were added and cells were incubated overnight at 4° C. Cells were then washed and incubated with fluorochrome conjugated secondary antibodies (Invitrogen, #A-11001A and # A-21428), diluted in blocking buffer for 1 hour at room temperature. The cells were washed and stained with DAPI for 10 minutes at room temperature in the dark. After a final wash, cells were stored in PBS at 4^°^ C prior to acquisition using the 100X oil immersion lens of a Nikon A1 confocal microscope (Nikon). For colocalization quantification, a python script based on NumPy, SciPy, pandas, and Scikit-image was used. Briefly, cell masks were used to define single cell regions of interest from which pixel intensities were extracted for each channel. Cells containing fewer than 20 pixels were excluded. Pearson’s correlation coefficient was calculated between AF488 and AF555 intensities within each cell when both channels exhibited non-zero variance. Manders overlap coefficients (M1 and M2) were computed as the fraction of AF488 signal overlapping AF555-positive pixels, and the fraction of AF555 signal overlapping AF488-positive pixels respectively. Differences in colocalization were determined by the Wilcoxon rank-sum test.

**Supplementary figures.**

**S1.** **Loading of BMDCs with homologous MHC epitopes skews differentiation towards a cDC1-like phenotype.**

**A.** Table summarizing statistical analysis of scRNA-seq cluster distribution between experimental conditions. N=3 mice. **B**. Heatmap showing the top differentially expressed genes between cluster 1 vs cluster 0. Colors represent scaled gene expression levels across cells (red, highest expression; blue, lowest expression). **C**. UMAP feature plot showing expression pattern of lineage-associated transcription factors *Mafb*, *Id2*, *Batf3*, and *Zeb2* used to define cluster 6 as the root for pseudotime inference. **D**. Expression pattern of selected genes associated with pseudotime progression along the Monocle inferred branching trajectory between cluster 1 and cluster 0. **E.** UMAP showing expression of select lineage-defining markers along the pseudotime trajectory. **F**. Table listing all genes associated with branching pseudotime progression identified during trajectory inference analysis.

**S2.** **cDC1-like cluster 1 is enriched for type 1 immune polarizing gene programs.**

**A.** GSEA plot showing the enrichment of antigen processing and proteasomal degradation pathway genes in cluster 1 compared cluster 0. **B.** Dot plot showing the expression of leading-edge genes driving the positive enrichment of antigen processing and proteasomal degradation pathway in cluster 1 compared cluster 0. Dot size represents the proportion of cells expressing each gene and color intensity represents scaled gene expression. **C.** GSEA plot showing positive enrichment of antigen processing and presentation on MHC II pathway genes in cluster 1 compared to cluster 0. **D.** Dot plot showing the expression of leading-edge genes contributing to the positive enrichment of antigen processing and presentation on MHC II pathway in cluster 1. **E.** GSEA plot showing the enrichment of IL2-Stat5 signaling pathway genes in cluster 1 compared to cluster 0. **F.** Dot plot showing the expression of leading-edge genes contributing to the positive enrichment of IL2-Stat5 signaling pathway in cluster 1. **G.** GSEA plot showing the negative enrichment of IL6-Jak-Stat3 signaling pathway genes in cluster 1. **H.** Dot plot showing the expression of leading-edge genes driving the negative enrichment of IL6-Jak-Stat3 signaling pathway in cluster 1.

**S3.** **mTORC1 pathway signaling is downregulated in the cDC1-like cluster**

**A**. Heatmap showing inferred signaling pathway activity across scRNA-seq clusters using PROGENy pathway analysis. **B**. Heatmap showing the expression of leading-edge genes driving the negative enrichment of mTORC1 signaling pathway in cluster 1 relative to all other clusters. **C**. Heatmap of reverse phase protein array (RPPA) data showing levels of total and phosphorylated proteins. N=4 mice.

**S4.** **Type 1 immune polarization is driven by NF-κB-mediated transcriptional reprogramming downstream of mTORC1 inhibition.**

Transcription factor binding motifs enriched within NF-κB peaks in the BMDC loaded with heterologous control (**A**) or homologous MHC peptides (**B**) as determined by Homer motif analysis. **C.** Representative genome browser tracks (mm10 annotation) showing NF-κB ChIP-seq occupancy at *Il12b* gene. N=3 mice. **D.** Validation of *IL12b* expression in human moDC. Data represents n=6 donor samples pooled from 3 independent experiments. Statistical analysis performed by paired t test. **P<0.01.

**S5.** **Inhibition of mTOR enhances the cDC1-like phenotype and drives DC-mediated type 1 immune responses in T cells**. **A-B.** Experimental workflow for ex vivo inhibition of mTORC1 signaling in BMDC. BMDC were differentiated from C57/BL6 mice bone marrow and loaded with either homologous or heterologous MHC peptides. After 3-hour incubation with the peptides, mTORC1 signaling was inhibited for 6h. Rapamycin was subsequently washed out and cells cultured with complete media supplemented with maturation cocktail. After 24h, DCs were analyzed by flow cytometry, and the supernatant was processed for Il12 and Il10 ELISA. **A.** Representative FACs plots displaying CD80 expression in BMDC loaded with MHC peptides with or without rapamycin treatment, with corresponding quantification. N= 7 mice. **B.** Representative FACs plots displaying CD86 expression in BMDC loaded with MHC peptides with or without rapamycin treatment, with corresponding quantification. N=4 mice. **C-F**. Experimental workflow for ex vivo stimulation of human PBMC by DC with or without rapamycin treatment. Monocytes from healthy donors were differentiated into moDC. Immature moDC were loaded with MHC peptides and treated with rapamycin for 6h to inhibit mTOR signaling, followed by maturation for 48h. Dendritic cells were then co-cultured with allogeneic PBMCs from healthy donors and restimulated on day 9. After 14 days, T cell subsets were characterized by flow cytometry. **C**. Representative FACs plots displaying Tbet+ CD8+ T cells in the DC-PDMC co-culture, with corresponding quantification. N=5 healthy donors. **D**. Representative FACs plots displaying PD1+ CD8+ T cells in the DC-PBMC co-culture, with corresponding quantification. N=4 healthy donors. **E**. Representative FACs plots displaying PD1+ CD4+ T cells in the DC-PBMC co-culture, with corresponding quantification. N=4 healthy human donors. **F.** Representative FACs plots displaying GATA3+ CD4+ T cells in DC-PBMC co-culture, with corresponding quantification. N=4 healthy donors. Statistical analysis was performed by two-way ANOVA followed by Sidak’s multiple comparison test. p < 0.05, ∗∗p < 0.01, and ∗∗∗p < 0.001

**S6. The cluster 1 cDC1-like gene signature is enriched among human clinical trial moDC vaccine products**. **A.** Heatmap showing expression of leading-edge genes driving the positive enrichment of the mouse cDC1-like cluster in homologous antigen-loaded human moDC clinical vaccine products compared to the control.

**Table S1.** List of antibodies and probs used in the study.

| **List of antibodies for flow cytometry analysis** | | | | | | |
| --- | --- | --- | --- | --- | --- | --- |
| **Target** | **Conjugate** | **Clone** | **Dilution** | **Source** | **Catalog#** | **RRID** |
| CD3 | APC | SK7 | 1:25 | Biolegend | 344812 | AB_10645473 |
| CD4 | PECY5 | RPA-T4 | 1:25 | Biolegend | 300510 | AB_314078 |
| CD8 | BUV737 | SK1 | 1:25 | BD Horizon™ | 612755 | AB_2870086 |
| CD25 | AF488 | M-A251 | 1:25 | Biolegend | 102017 | AB_2870086 |
| PD-1 | PE | NAT105 | 1:25 | Biolegend | 367404 | AB_2566065 |
| IFNg | BV711 | 4S.B3 | 1: 25 | BD Horizon™ | 564793 | AB_2738953 |
| GRANZYME B | PECY7 | QA18A28 | 1: 25 | Biolegend | 396410 | AB_280107 |
| Tbet | BV605 | 4B10 | 1: 25 | Biolegend | 644817 | AB_11219388 |
| GATA3 | APC-Fire | W19195B | 1: 25 | Biolegend | 386906 | AB_3097419 |
| LIVE CELLS | Zombie UV |  | 1:500 | Biolegend | 423108 |  |
| XCR1 | PE | ZET | 1: 25 | Biolegend | 148204 | AB_2563843 |
| CD11b | UV395 | M1/70 | 1:25 | Biolegend | 101240 | AB_3097646 |
| CD11c | PECY7 | N418 | 1:25 | Biolegend | 117318 | AB_493568 |
| IA-IE | FITC | M5/114.15.2 | 1:25 | Biolegend | 107606 | AB_313321 |
| SIRPa | BUV737 | P84 | 1:50 | BD Horizon™ | 741819 | AB_2871154 |
| BST2 | BV480 | 927 | 1:25 | BD Horizon™ | 747600 | AB_2744168 |
| SIGLEC-H | AF408 | E50-2440 | 1:25 | BD Horizon™ | 567005 | AB_2870001 |
| CD86 | BV650 | GL-1 | 1:50 | Biolegend | 105036 | AB_2686973 |
| CD80 | APC | 16-10A1 | 1:50 | Biolegend | 104714 | AB_313135 |
| **List of antibodies for immunofluorescence staining** | | | | | | |
| **Target** | **Conjugate** | **Clone** | **Dilution** | **Source** | **Catalog#** | **RRID** |
| mTOR | Unconjugated |  | 1:300 | Cell Signaling Technology | 2983 | AB_2105622 |
| LAMP2 | Unconjugated |  | 1:125 | Abcam | ab25631 | AB_470709 |
| Goat anti-Rabbit  IgG | AF555 |  | 1:400 | Invitrogen | A-21429 | AB_2535850 |
| Goat anti-Mouse IgG | AF647 |  | 1:400 | Invitrogen | A-21235 | AB_141693 |
| Donkey Anti-Mouse IgG | AF647 |  | 1:300 | Jackson Immunoresearh Laboratories | 715-605-151 |  |
| Donkey Anti-Rabbit IgG | AF568 |  | 1:300 | Jackson Immunoresearh Laboratories | 711-575-152 |  |
| List of antibodies for immunoblotting | | | | | | |
| **Target** | **Conjugate** | **Clone** | **Dilution** | **Source** | **Catalog#** | **RRID** |
| p-mTOR  (Ser2448) | Unconjugated |  | 1:1000 | Cell Signaling Technology | 2971 | AB_3668749 |
| mTOR | Unconjugated | 7C10 | 1:1000 | Cell Signaling Technology | 2983 | AB_2105622 |
| p-p70S6K  (Thr389) | Unconjugated |  | 1:1000 | Cell Signaling Technology | 9205 | AB_10078518 |
| P70S6K | Unconjugated | 49D7 | 1:1000 | Cell Signaling Technology | 5707 | AB_10694087 |
| P-cFos  (Ser32) | Unconjugated | D82C12 | 1:1000 | Cell Signaling Technology | 44436 | AB_2897355 |
| c-Fos | Unconjugated | 9F6 | 1:1000 | Cell Signaling Technology | 2250 | AB_3086834 |
| beta-Actin | Unconjugated | 13E5 | 1:1000 | Cell Signaling Technology | 5125S | AB_1903890 |
| Anti-rabbit IgG | Unconjugated | polyclonal | 1:1000 | Cell Signaling Technology | 7074 | AB_3718177 |
| Probes used for qPCR | | | | | | |
| Target | Probe | Source |  |  | Catalog# | RRID |
| IL12a | Hs01076226_m1 | Invitrogen |  |  | 4351372 |  |
| IL10 | Hs00961622_m1 | Invitrogen |  |  | 4331182 |  |
